# Leveraging Human Pangenome for Improved Somatic Variant Detection

**DOI:** 10.64898/2026.01.04.697580

**Authors:** Qichen Fu, Zilan Xin, Benpeng Miao, Wenjin Zhang, Nahyun Kong, Zitian Tang, Andrew Ruttenberg, Derek Albracht, Edward A. Belter, John E. Garza, Chad Tomlinson, Elvisa Mehinovic, Jiawei Shen, Xiaoyu Zhuo, Shihua Dong, Benjamin K. Johnson, Mary F. Majewski, Theron Palmer, H. Josh Jang, Yuchen Cheng, Zefan Li, Heather A. Lawson, Tina Lindsay, Daofeng Li, Robert Fulton, Hui Shen, Sheng Chih Jin, SMaHT Network Assembly/Pangenome Working Group, Juan F. Macias-Velasco, Ting Wang

## Abstract

Somatic variant detection is technically challenging due to low variant allele fractions, the confounding presence of germline variation, and reference bias. Linear references such as GRCh38 miss sample-specific variation, causing misalignments and incorrect variant calls. Although telomere-to-telomere donor-specific assemblies (DSAs) accurately represent individual genomes, their application is limited by cost and technical barriers. Alternatively, the graph-based human pangenome provides a scalable framework to improve read alignment and perform genome inference. Here, we benchmarked somatic variant detection using GRCh38, graph-based pangenomes, and pangenome-inferred DSAs with a HapMap mixture dataset and the COLO829 melanoma cell line. Pangenome-guided alignment improves read mapping and somatic variant calling accuracy. Furthermore, personalized pangenomes partially reconstruct donor-specific genomic content, improving accuracy, reducing germline contamination, and enabling detection of events in loci absent or poorly represented in GRCh38. These findings demonstrate that graph-based and personalized pangenomes are effective strategies for enhancing somatic variant detection compared with GRCh38.

## Introduction

Somatic variants arise after fertilization and accumulate throughout an individual’s lifetime.^1^ They are primarily caused by errors in DNA replication or exposure to exogenous DNA-damaging agents, such as ultraviolet radiation, chemicals, X-rays, and tobacco smoke.^2–5^ If left unrepaired or not being repaired correctly, these DNA lesions can result in base substitutions, small insertions or deletions, or large structural changes.^3^

While most somatic mutations are silent with minimal phenotypic consequences, a subset contributes to cancer, aging, and neurological disorders.^6^ However, the exact impact of somatic mosaicism on biological function remains poorly understood.^2,4,5^ The Somatic Mosaicism Across Human Tissues (SMaHT) network aims to shed light on how these events arise and how they impact human health and disease.^2^ This will be achieved by generating a comprehensive and reliable catalogue of somatic variation across 19 non-diseased tissue types from 150 donors.^2^

A major barrier for this endeavor is the technical challenge of somatic variant detection. Unlike germline variants, somatic mutations typically occur in a small fraction of cells and exhibit considerable heterogeneity across cells, tissues, and individuals. Low variant allele fractions (VAFs), combined with sequencing errors and artifacts introduced during library preparation, make accurate detection particularly difficult. Moreover, when alignment-based methods are used, the accuracy of somatic variant detection relies on the reference genome. GRCh38 is a mosaic haploid genome composed of a small number of individuals.^7,8^ It remains incomplete, with 6.7% of primary chromosome scaffolds unresolved, and its ancestry representation is limited, comprising approximately 57% European and 37% African ancestry.^7,8^ Traditional somatic variant detection methods based on GRCh38 rely on either paired normal-tumor samples or panel-of-normal strategies to filter out germline variants. These approaches are still vulnerable to biases introduced using an under-representative reference genome.^7,9,10^ The genetic distance between GRCh38 and individual genomes creates reference biases that affect read mapping rate and accuracy, which leads to increased incorrect variant calls. For somatic variant calling, reference biases increase germline contamination in the variant call set, leading to higher chances of misclassification between germline and somatic variants^10^. Moreover, somatic variants located in regions absent from GRCh38 are undetectable when using GRCh38 as the reference.^7^

Because no single linear reference genome can adequately capture human genomic diversity, alternative strategies are needed. While high-quality, donor-specific assemblies (DSAs) are optimal references for somatic variant detection, their high cost limits broad adoption. To overcome these limitations, the Human Pangenome Reference Consortium (HPRC) proposed a shift from a fixed linear reference to a pangenome: a genome-graph composed of a collection of individuals that are representative of most of the common genetic variation from diverse populations.^11^ The HPRC is assembling genomes from 350 individuals of diverse ancestries, and in 2023, released a draft human pangenome reference consisting of 94 assemblies from 47 samples and a genome graph incorporating 90 haplotypes, anchored by both GRCh38 and T2T-CHM13.^12^ This resource has already demonstrated improved detection of single nucleotide variants (SNVs) in short-read sequencing data.^12^

In this study, we presented two strategies to apply pangenome to somatic variant detection: 1) alignment to the pangenome itself 2) use of a pangenome to approximate the germline assembly of the individual to be analyzed. We benchmarked somatic variant calling across three reference types: GRCh38, pangenome graphs, and pangenome-inferred DSAs. Benchmarking was performed using two experimental systems: a HapMap cell line mixture (Figure 1A) and the COLO829 melanoma cell line (Figure 1B). SMaHT network designed and created the HapMap cell line mixture—a mixture of cell lines in which HG005 contributes 83.5% of the mixture (corresponding to germline variants) and five other donors contribute 0.5-10% of the mixture (corresponding to somatic variants) (Figure 1A).^13^ The COLO829 melanoma cell line and its matched normal B lymphoblast cell line (COLO829BL) were also used for benchmarking (Figure 1B). We utilize short-read and long-read whole-genome sequencing (WGS) data of two experimental systems and the HapMap mixture benchmarking variant sets^14^ to compare alignment metrics, precision/recall of somatic variant calling (SNVs, indels, and SVs), and mutational signatures (single-base substitution (SBS) & double-base substitution (DBS)) using different references (Figure 1C and 1D).

**Figure 1.**
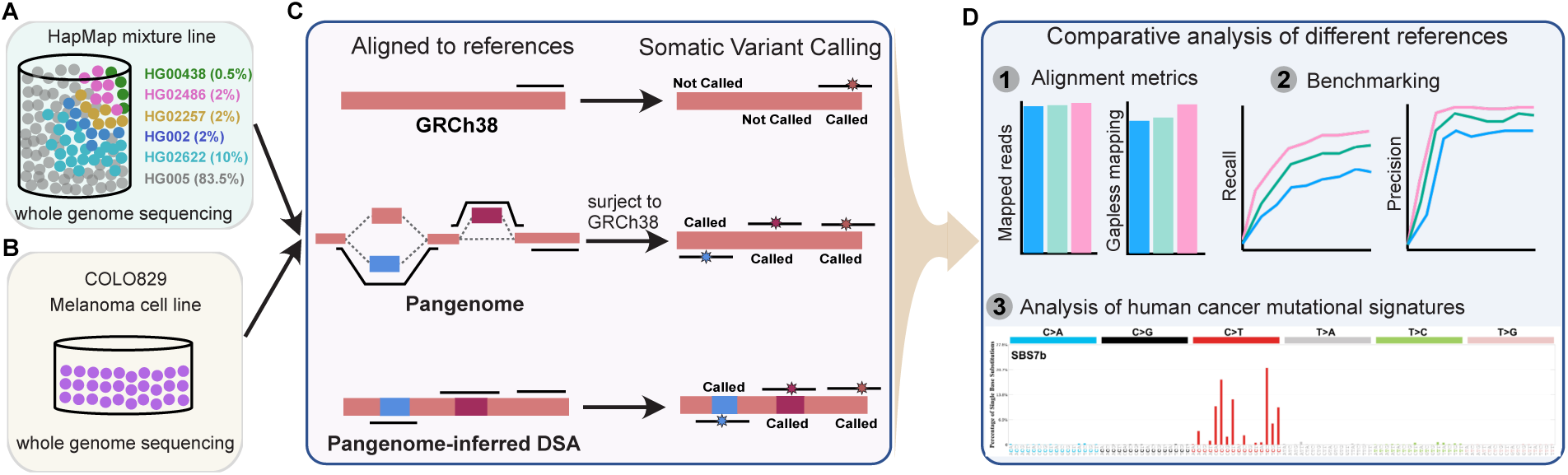
Schematic workflow of the benchmark study. **A.** Benchmark datasets, including the HapMap mixture (500x Illumina short reads, 100x PacBio HiFi long reads), which was generated by pooling six HapMap cell lines composed of 83.5% HG005 representing the germline background and five additional HapMap donors contributing at 0.5–10% each to simulate somatic variation across a spectrum of VAFs, **B.** and the COLO829 melanoma cell line with its paired normal control (COLO829BL) providing a cancer-derived context. **C.** Overview of the reference types evaluated in this study, including GRCh38, graph-based pangenomes, and pangenome-inferred DSAs. **D.** Alignment of sequencing reads to each reference type using either linear or graph-based aligners, followed by variant calling of SNVs, small indels, and structural variants with appropriate tools. Benchmarking strategy in which variant calls were compared to mixture-derived benchmarking variant sets or tumor–normal comparisons, with stratification by genomic region difficulty and variant allele fraction. Downstream analyses of alignment metrics, variant calling precision and recall, and mutational signature profiling to assess the impact of reference choice on somatic variant detection accuracy.

Our analyses showed that alignment to the pangenome improves alignment quality for short-read and long-read sequencing and subsequently increases the accuracy of somatic variant calling. Pangenome personalization can partially approximate a DSA, which further improves variant calling accuracy, enables better classification between somatic and germline variants, and allows for more comprehensive somatic variant profiling.

## Results

### Pangenome represents genetic diversity and improves the alignment quality for short and long reads

To benchmark pangenome-based somatic variant detection against linear reference (GRCh38), we utilized HapMap cell line mixture composed of 83.5% HG005 (Male; Chinese) and 10% of HG02622 (Female; Gambian in Western Division, Mandinka), 2% each of HG002 (Male; Ashkenazim Jewish), HG02257 (Female; African Caribbean in Barbados), and HG02486 (Male; African Caribbean in Barbados), and 0.5% of HG00438 (Female; Chinese Han in the South) (Figure 1A)^12^. The variants present in HG005 but absent in any other five individuals are considered as germline variants, whereas the variants present in any of the five individuals but absent in HG005 are considered as somatic variants.

To evaluate whether the pangenome can represent HapMap germline and somatic samples without their assemblies existing in the graph, we first generated two pangenome graphs using Minigraph-Cactus^15^ and compared their nodes, edges, and degree of edges with the HPRC v1.1 pangenome^12^. The HPRC v1.1 pangenome (Graph v1 in Figure 2A), constructed by Minigraph-Cactus^15^, contains GRCh38, CHM13, and 44 diploid assemblies from diverse genetic backgrounds (including four of the six HapMap samples: HG00438, HG02257, HG02486, HG02622, and excluding HG002 and HG005). The v2 graph contains all HPRC Release 1 assemblies, including HG005, but excluding the five somatic samples. The v3 graph excludes all six HapMap samples (Figure 2A). Three pangenome graphs show a highly similar number of nodes and edges (Figure 2B). The degree (the number of connections to the node) distributions of nodes confirm that the overall graph complexity properties are comparable across versions (Figure 2C). The majority of variants in HapMap samples are captured by other individuals in the v3 pangenome graph, indicating the pangenome graph without HapMap samples can be recognized as a representative reference (Figure S1).

**Figure 2.**
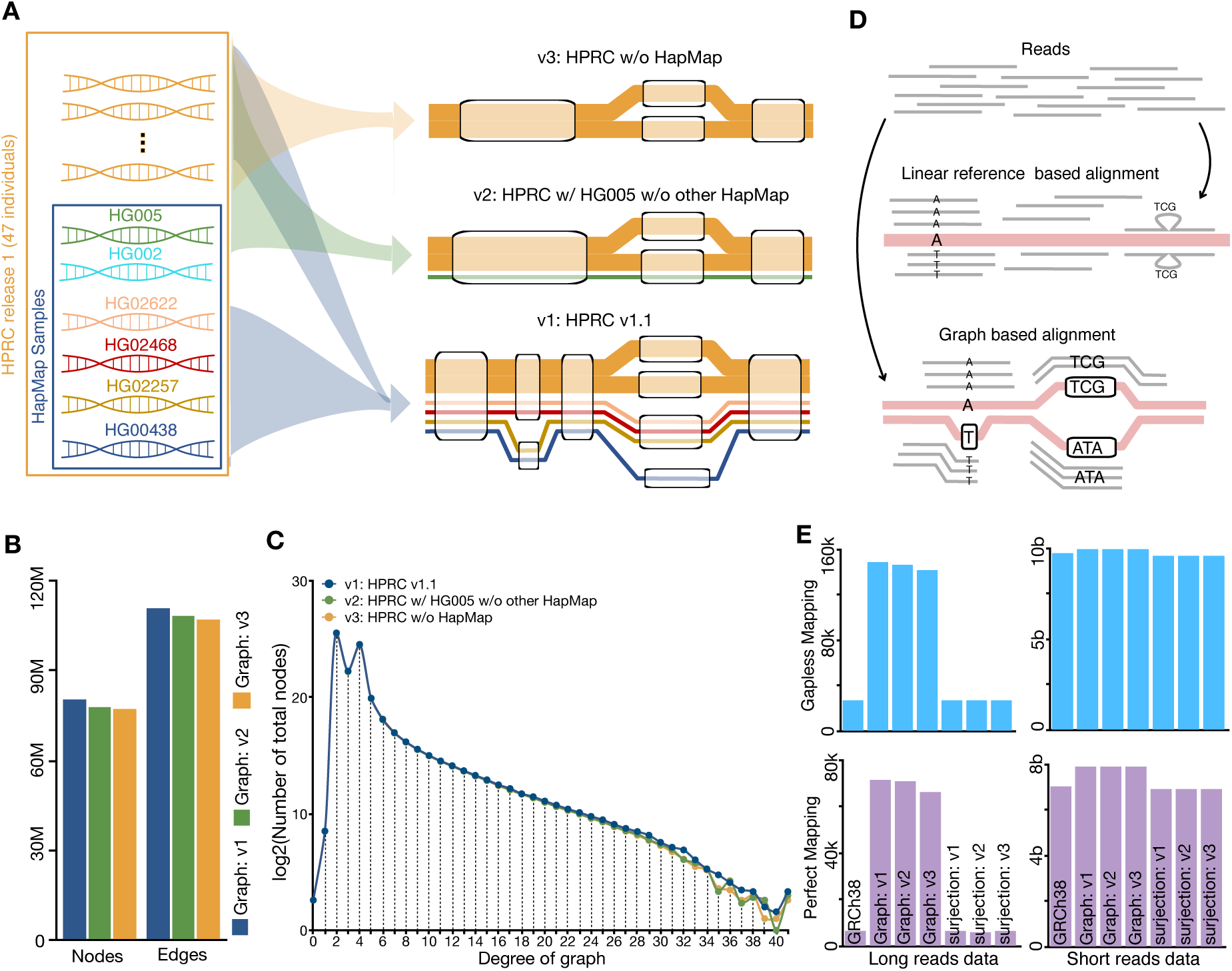
Evaluation of how pangenome composition influences variant representation and alignment quality. Three versions of the draft human pangenome graph were constructed to test whether variants from HapMap mixture donors require explicit inclusion or are already captured through sharing with other assemblies. **A.** Graph v1 includes GRCh38, CHM13, and 44 diploid assemblies (including four HapMap donors). Graph v2 includes all HPRC Release 1 assemblies with HG005 but excludes the five other HapMap donors. Graph v3 excludes all six HapMap donors. **B.** Node and edge counts for the three graphs show highly similar structures, indicating that most variants from the HapMap donors are represented on other assemblies in the pangenome. **C.** Degree distributions confirm that the overall graph complexity properties are comparable across versions. **D.** Illumina short reads (500x) and PacBio HiFi long reads (100x) from the HapMap mixture were aligned to each graph using vg giraffe, resulting in more perfectly mapped and gapless reads than alignment to GRCh38. **E.** Alignment quality metrics comparing graph-based alignments with GRCh38 show that the pangenome improves read placement and alignment score. When graph alignments are surjected back to GRCh38, the proportion of perfectly mapped and gapless reads decreases. In this context, however, the decrease does not indicate poorer alignment quality but instead reflects increased sensitivity to real sequence divergence, such as small indels, that are captured during graph alignment and retained after surjection.

To determine whether alignment to the pangenome can improve alignment quality, 500x Illumina short-read and 100x PacBio HiFi long-read were aligned to three pangenome graphs (clipped and allele filtered) using vg giraffe^16^ (Figure 2D). All pangenome graphs achieved slightly lower numbers of mapped reads (Figure S2). However, the alignment quality was substantially improved with a notable and comparable increase in the number of properly paired, gapless, and perfectly mapped reads for both long and short reads compared with GRCh38 (Figure 2E and S2). For short reads, the gapless and perfectly mapped reads increase by 1.76% and 12.46% respectively (Figure 2E). For long reads, the gapless and perfectly mapped reads increase by 425.07% and 966.27% respectively (Figure 2E). To be compatible with linear variant callers, graph alignments were converted to linear (GRCh38) alignments through surjection. Surjection partitions an alignment: the portion aligned to reference nodes remains unchanged, while the portion aligned to non-reference nodes is projected onto nearby reference nodes via local alignment.^12^ This process decreases the number of perfectly mapped and gapless reads. However, the pangenome anchors reads in accurate regions and generates more even coverage across the genome compared to GRCh38 (Figure S3). These results indicate that using a pangenome as a more representative reference improves alignment quality compared with GRCh38, substantiating previous results^12,17^.

### Pangenome improves somatic variant calling accuracy through improved alignment quality

To understand whether the improved alignment quality can improve somatic variant calling accuracy, we compared the variant calling precision and recall from pangenome-guided alignment with GRCh38 alignment. The short-read and long-read WGS data of the HapMap mixture were aligned to the v3 pangenome graph (clipped and allele filtered) with alignments surjected to GRCh38. Small variants (SNVs & indels <50 bp) were called and filtered from Illumina short read alignments using MuTect2^18^. SVs were called and filtered from PacBio HiFi long reads using Sniffles2^19^. To benchmark the variant calling accuracy, we utilized benchmarking variant sets generated using a genome-graph approach established by Kong et al^14^, with GRCh38 as references (Table S1). The pangenome improves SNV calling precision and recall across all VAF bins between 0 and 16.5% (with overall precision improving from 0.9625 to 0.9734 and recall from 0.7145 to 0.7284) (Figure 3A and Table S2). The improvement is greatest in extreme regions, with SNV precision improving from 0.7908 to 0.8506 and recall from 0.4802 to 0.5069 (Figure 3A and Table S2). Moreover, no evident differences in precision or recall are observed across the v1, v2, and v3 pangenome graphs (Figure S4), suggesting that the pangenome can improve variant calling accuracy regardless of whether a DSA is present in the graph.

**Figure 3.**
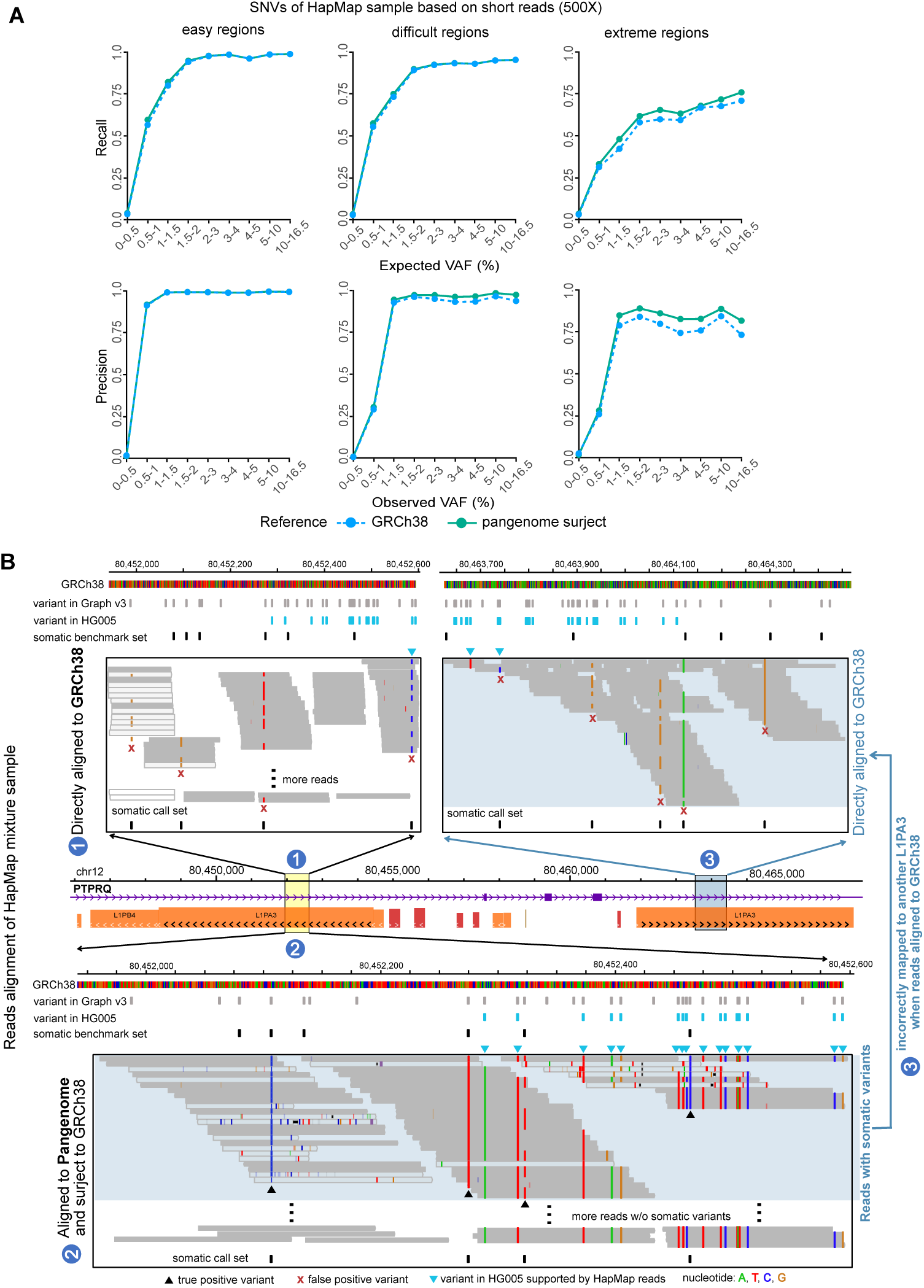
Pangenome improves somatic variant detection in the HapMap mixture. **A.** Precision and recall for SNVs across VAF bins from 0-16.5%, showing consistent improvements in both measures when using the pangenome, with the largest gains in extreme regions. **B.** Example locus on chromosome 12 where a somatic SNV is detected when aligning reads to the pangenome but not when aligning to GRCh38. Supporting reads are present in both cases; however, under GRCh38 alignment, the reads are misplaced to a different locus, resulting in an apparent false negative. Benchmarking variant set information confirms the variant at the pangenome-supported locus, illustrating how the pangenome improves correct read placement and reduces false negative and false positive calls. Comparison of read placement for the same supporting reads under GRCh38 alignment, showing their misalignment to a different genomic region.

As an example of how pangenome alignment reduces false negative and false positive somatic variant calls, we highlight a locus at chr12:80,449,114-80,466,689, which contains two repetitive L1PA3 elements (Figure 3B). The somatic variants were detected when reads were aligned to the pangenome but not when aligned to GRCh38. The supporting reads are present in both cases; however, under GRCh38 alignment, they are misplaced to a different genomic region, resulting in apparent false negatives. Incorporating the benchmarking variant set information confirms that the variant resides at the pangenome-supported locus, illustrating how the pangenome improves read localization and enables more accurate somatic variant detection. As indicated in the pangenome deconstructed VCF^12^, the two regions have more non-reference sequences, suggesting that the pangenome increases mappability by introducing germline variants that allow aligners to distinguish highly repetitive elements. Graph-based aligners can leverage these non-reference sequences to more effectively localize read placement among highly similar sites.

Alignment to all v1, v2, and v3 pangenome graphs improves somatic indel calling precision (from 0.7410 to 0.7564) and leads to comparable recall between alignment to GRCh38 (0.1867) and pangenome (0.1860) (Figure S5A and Table S2). Precision and recall for SVs are comparable between GRCh38 (0.6640; 0.2024) and pangenome (0.6598; 0.1939) (Figure S5B and Table S2). Moreover, PacBio HiFi long reads span much longer ranges than Illumina short reads, making them less difficult to align accurately on GRCh38. By comparing reads aligned to GRCh38 with those aligned to the pangenome and then surjected to GRCh38, we found that a larger proportion of short reads were mapped to distinct genomic locations between the two approaches compared to long reads (Figure S6), suggesting pangenome benefits less with long read alignment than short read alignment. Together, these results suggest that the pangenome can improve somatic variant calling accuracy through improved alignment quality, especially in challenging genomic regions.

### Pangenome enables genome inference, improves somatic variant calling accuracy, and reduces germline contamination

Surjecting reads from the pangenome onto GRCh38 utilizes improved alignments derived from the pangenome representativeness but does not fully take advantage of the completeness and genetic diversity of the pangenome, including sequence that exists in individuals but are missed in GRCh38. Next, we used the pangenome to infer and partially reconstruct the germline assembly.

Variants that are in pangenome graphs but not part of the sample will mislead read mappings. To address this, pangenome personalization has been developed as a strategy to remove irrelevant variants and impute a personalized pangenome subgraph by sampling local haplotypes according to k-mer counts in the individual’s sequencing data.^17^ This approach further improves read mappings, variant calling, and genotyping.^17^ This also allows for leveraging an individual’s sequencing data to construct artificial haplotypes that approximate the person’s genomic background, thereby enabling the imputation of a DSA.

Here, we used the personalization approach (vg haplotype) to sample the haplotypes that match closest to the k-mer content of the sample from an allele-unfiltered pangenome graph and construct a subgraph containing pseudo-diploid assemblies (hap1 and hap2).^17^ The v3 pangenome graph (allele unfiltered, clipped, and excludes all 6 HapMap samples) and k-mer content of HG005 short read WGS were used as an input for pangenome personalization (Methods). To determine how closely inferred DSAs match a high-quality DSA, we aligned GRCh38 and the pangenome-inferred DSAs of HG005 hap1 and hap2 with HG005 diploid assemblies using Minimap2^20^. The pangenome-inferred DSA shows lower genetic divergence with HG005 than GRCh38, with fewer substitutions, insertions, and deletions (Figure 4A). To tolerate haplotype switching between pangenome-inferred DSA hap1 and hap2 in pangenome personalization, we also evaluated the pangenome-inferred DSAs using a graph-based approach (Figure 4B). Both the pangenome-inferred DSAs (hap1 and hap2) and the HG005 diploid assembly were incorporated into a Minigraph-Cactus^15^ validation graph. To compare genotypes between the pangenome-inferred assembly and HG005, the validation graph was deconstructed to a VCF with genotype information for four assemblies. Variants in HG005 paternal or maternal assembly were considered captured if they were present in either pangenome-inferred DSA hap1 or hap2. Missing variants were caused by assembly gaps in the HG005 maternal or paternal assembly (Figure 4B). The pangenome-inferred DSAs were found to capture most HG005 variants (Maternal: 84.7% captured, 6.88% missing, 8.38% not captured; Paternal: 86.8% captured, 6.88% missing, 6.33% not captured) (Figure 4C).

**Figure 4.**
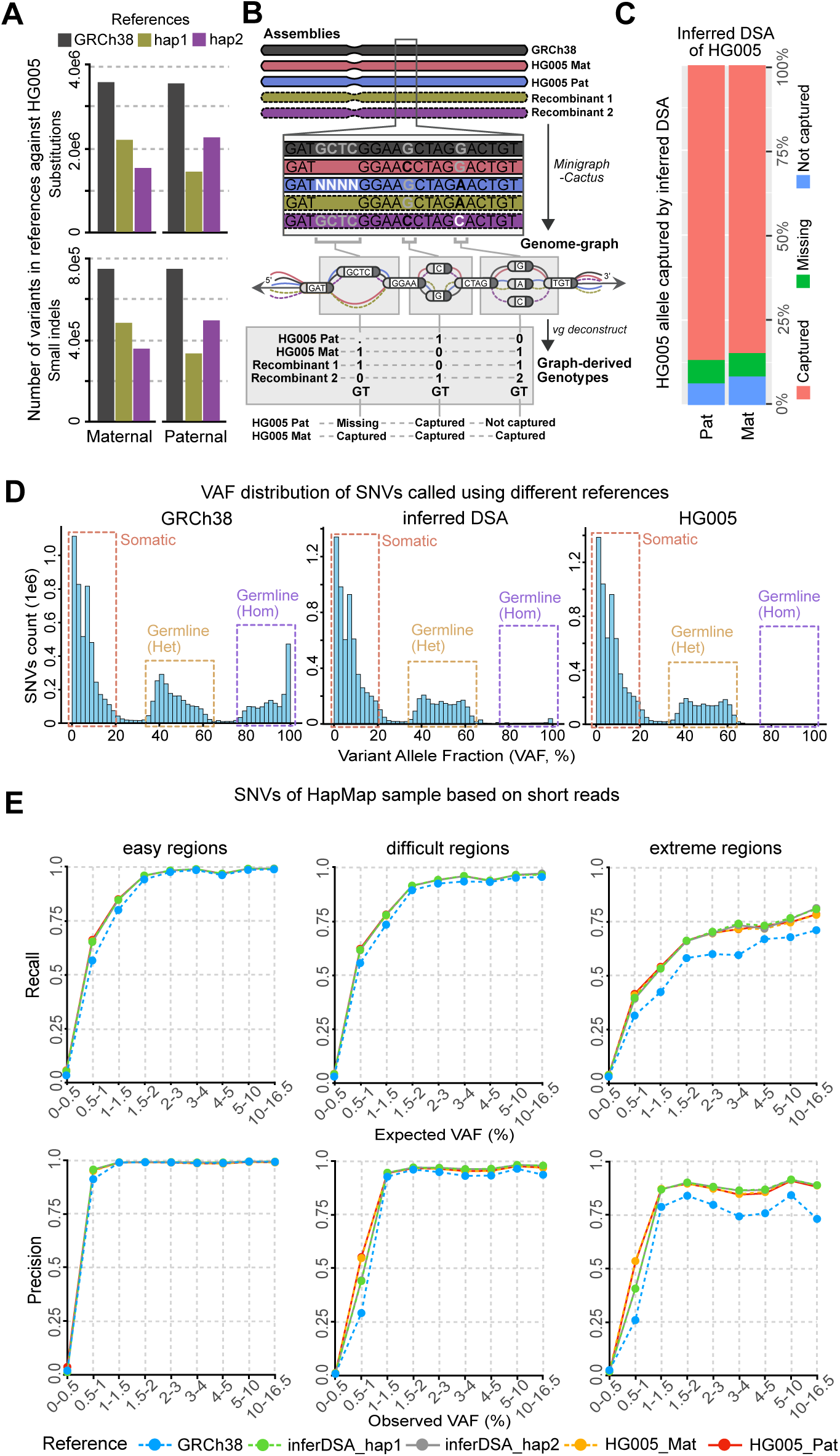
Pangenome-inferred assemblies improve somatic variant detection in the HapMap mixture. Construction of a pangenome-inferred assembly for HG005 using k-mer content from short-read data and haplotype sampling from the v3 pangenome graph, which excluded all 6 HapMap samples. The resulting pseudo-diploid assembly partially reconstructs the HG005 genome. **A.** Comparison of sequence divergence between the inferred HG005 assembly, the true HG005 maternal and paternal assemblies, and GRCh38, showing that the inferred assembly reduces the number of substitutions relative to GRCh38. **B.** A graph-based method was employed to jointly compare genotypes between the pangenome-inferred assembly and HG005. **C.** Graph-based genotype comparisons show that pangenome-inferred assembly captures approximately 90% of HG005 variants, demonstrating its ability to approximate a donor-specific assembly. **D.** VAF distributions of non-VAF filtered SNV call sets from MuTect2 tumor-only mode for GRCh38, pangenome-inferred DSA (hap1), and HG005 paternal assembly. **E.** Benchmarking of somatic variant calling performance across reference types. The pangenome-inferred DSA achieves precision and recall for SNVs comparable to the HG005 maternal and paternal assemblies and superior to GRCh38, with the largest gains observed in difficult and extreme genomic regions.

Given the improvement in representation from GRCh38 to pangenome-inferred DSA, we next benchmarked somatic variant calling on GRCh38, pangenome-inferred DSAs, HG005 maternal, and HG005 paternal assemblies. Classifying germline and somatic variants becomes challenging when a matched normal is unavailable or inapplicable, such as in normal tissue^21^. The pangenome-inferred DSA filters the majority of HG005 homozygous germline variants at the alignment step (Figure 4D). The VAF distribution of the pangenome-inferred DSA approximates that of the HG005 assemblies (Figure 4D). To benchmark somatic variant calling accuracy, we utilized benchmarking variant sets generated with a genome-graph approach established by Kong et al.^14^, with the pangenome-inferred DSAs (hap1 and hap2) and the HG005 maternal and paternal assemblies as references (Figure S7 and Table S1). These sets captured all possible variants across SNVs, indels, and SVs and were used to compare with respective call sets. The pangenome-inferred DSA hap1 and hap2 achieved precision and recall comparable to the HG005 maternal and paternal assemblies for SNVs, indels, and SVs, and exceeded those of GRCh38 (Figure 4E, S8, and Table S3). The improvement was most pronounced in difficult and extreme regions (Figure 4E, S8, and Table S3). The pangenome-inferred DSA outperforms pangenome-guided alignment in variant calling for SNVs, indels, and SVs (with SNV precision improving from 0.9734 to 0.9764 and recall from 0.7284 to 0.7819 genome-wide; precision from 0.8506 to 0.8871 and recall from 0.5069 to 0.5912 in extreme regions) (Table S2 and S3).

As an example of how pangenome-inferred DSA reduces germline contamination and improves somatic variant calling accuracy, we looked at the same region in Figure 3B. We found that pangenome-inferred DSA correctly identified variants in the first L1PA3 element and filtered germline variants at the alignment step (Figure S9).

Together, these results show that pangenome-guided inference of donor-specific assemblies reduces germline contamination and enhances somatic variant detection to a level approaching that of true DSAs, while avoiding some of the limitations of surjection back to GRCh38.

### Pangenome-inferred DSA allows for more comprehensive somatic SNV profiling for COLO829

Since pangenome-inferred DSA partially reconstructs the germline assembly, we asked whether its call set captures a more comprehensive somatic SNV profile than the GRCh38 call set in the cancer sample. We utilized the PacBio HiFi data of the melanoma cell line COLO829 and its matched normal COLO829BL cell line (B lymphoblast cell line). We profiled the somatic SNVs of COLO829 using GRCh38 or pangenome-inferred DSA of COLO829BL using DeepSomatic^22^ tumor-normal mode. The pangenome-inferred DSA was constructed using HPRC v1.1 pangenome (allele unfiltered and clipped) and COLO829BL short-read WGS (20x)^17^. We validated the pangenome-inferred DSA by comparing it with the near T2T DSA of COLO829 generated using Verkko2^23^ by the UWSC Genome Characterization Center (GCC)^24^. The pangenome-inferred DSA captures most of the COLO829 germline variants (Maternal: 88.4% captured, 2.73% missing, 8.85% not captured; Paternal: 85.1% captured, 2.73% missing, 12.1% not captured) (Figure 5A). It improves the alignment quality with increased mapped reads and decreased insertion/deletion and supplementary alignments (Figure 5B). The pangenome-inferred DSA detects more somatic SNVs compared to GRCh38, including 4,290 SNVs unique to pangenome-inferred DSA and 930 SNVs unique to GRCh38 (Figure 5C).

**Figure 5.**
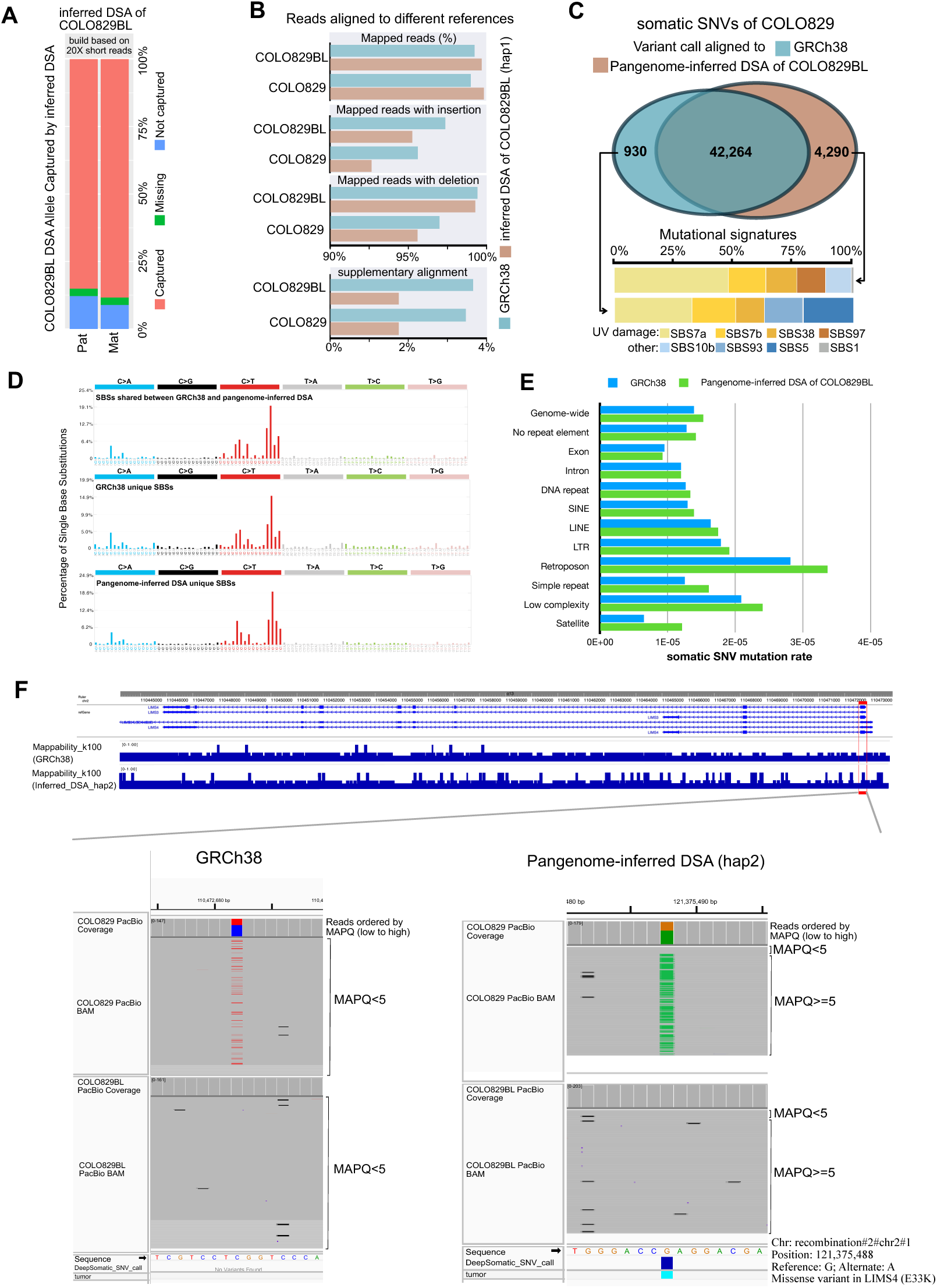
Pangenome-inferred DSA allows for more comprehensive somatic SNV profiling for COLO829. **A.** Allele capture rate of pangenome-inferred DSA of COLO829BL constructed based on 20x short-read WGS of COLO829BL. **B.** Alignment metrics of COLO829 and COLO829BL PacBio reads aligned to GRCh38 and pangenome-inferred DSA. **C.** The overlap between the SNV call sets using GRCh38 or inferred DSA as reference and the SBS signature decomposition of the unique SNVs. **D.** SBS profiles for shared and unique SNV calls of GRCh38 and pangenome-inferred DSA. **E.** Somatic SNV mutation rate of GRCh38 and pangenome-inferred DSA call set in different genomic features. Exon: total length of exons from protein-coding genes; Intron: total length of introns from protein-coding genes. **F.** Pangenome-inferred DSA improves the mappability and enables somatic variant detection in the LIMS4 gene.

Next, to assess the biological relevance of SNVs, we performed mutational signature analysis for shared and unique somatic SNVs from different references. The single-base substitution (SBS) and double-base substitution (DBS) signatures of shared SNVs were enriched in UV light exposure signatures (SBS7a, SBS7b, SBS38, and DBS1) (Figure S10 and S11). Unique SNV calls from pangenome-inferred DSA show higher SNV counts and proportion in UV light exposure SBS signatures (SBS7a, SBS7b, SBS38, and SBS97) than those of the unique SNV calls from GRCh38 (Figure 5C and 5D). DBSs unique to the pangenome-inferred DSA show a five-fold higher count in DBS1 compared with GRCh38-unique DBSs (Figure S10). The increased sensitivity in detecting mutations translated to a slightly higher estimated mutation rate when using pangenome-inferred DSA, both genome-wide and across major genomic features, except for satellite regions. Whereas using GRCh38 the estimated mutation rate in satellite regions was less than half the genome-wide rate, using the pangenome-inferred DSA doubled the estimated mutation rate in these regions (Figure 5E), highlighting that satellite regions are not less susceptible to UV damage and/or the DNA repair machinery is not more efficient^25,26^.

Among the 4,290 SNVs unique to the pangenome-inferred DSA, 2,070 of them can be lifted over to GRCh38. We observed that the pangenome-inferred DSA improves regional mappability, increases the MAPQ, and enables somatic variant detection. As an example, we highlighted the LIMS4 gene (Figure 5F). Across gnomAD^27^, GTEx^28^, and TCGA^29^, LIMS4 displays low population coverage, no detectable germline variants (Figure S12A), and low RNA expression across tissues (Figure S12B) and in melanoma (Figure S12C), yet elevated protein levels in melanoma (Figure S12C). This suggests that variants and RNA levels of the LIMS4 gene were underestimated due to the low mappability in GRCh38. We demonstrated that pangenome-inferred DSA increased the mappability and detected a missense mutation in the exon 1 of LIMS4, which was missed in the GRCh38 call set (Figure 5F). Moreover, the pangenome-inferred DSA aligned a greater number of somatic variant supporting reads, thereby enabling more sensitive and accurate somatic mutation detection (Figure S13).

A total of 2,220 pangenome-inferred DSA unique SNVs could not be lifted over to GRCh38. These SNVs are located in regions missing or poorly represented in GRCh38, including simple repeat expansion (Figure S14A), copy number gain (Figure S14B), and large missing segments, such as satellite regions (Figure 5E). Notably, the pangenome-inferred DSA determined that one haplotype of COLO829BL contains a simple repeat expansion within an intron of GNG10. This discovery is confirmed using COLO829BL DSA generated by UWSC-GCC^24^. A somatic SNV within the repeat expansion was detected using the pangenome-inferred DSA, which was missed by GRCh38 (Figure S14A). The pangenome-inferred DSA can also resolve copy number gains, resulting in improved coverage and more accurate somatic variant detection in extra copies, as illustrated by the example of LINC01410 (Figure S14B). We further summarized the pangenome-inferred DSA unique SNVs that fall within the exon (Table S4). 11 sSNVs were identified as missense mutations, 10 were in cancer genes, and 2 were predicted to be pathogenic by AlphaMissense^30^. 2 missense sSNVs were located in genes previously reported to be associated with melanoma, including ARHGEF5^31^, and TMEM87b^32^. 8 missense sSNV were in genes implicated in other cancer types, such as TRIOBP^33^, FCGBP^34^, and NBPF1^35^ (more in discussions and Table S4).

Together, these results highlight that the pangenome can construct a more representative assembly than GRCh38. This more representative assembly allows for more comprehensive and biologically meaningful somatic variant profiling.

## Discussions

The human pangenome has been developed to capture genetic variation in the human population^12^. It serves as a more complete and representative reference than GRCh38. Here, we presented the first comprehensive benchmark of somatic variant detection using pangenomic methods. We benchmarked pangenome-based somatic variant detection with the GRCh38 approach using two systems – the HapMap mixture and the COLO829 melanoma cell line, designed by the SMaHT Network. The benchmarking variant sets established by the genome-graph approach^14^ enable benchmarking of all variant classes, including SNVs, indels, and SVs. We demonstrated that the pangenome can capture most variants of the sample with sample’s assembly excluded and allows for improved alignment quality for both short and long read sequencing. This results in improvement in somatic SNV and indel calling, especially in challenging regions. Furthermore, the pangenome can partially reconstruct the germline assembly and improve the accuracy of somatic variant detection to a level comparable with the DSA. This also enables the more comprehensive somatic variant detection for cancer samples such as COLO829.

We demonstrated that the pangenome can improve alignment quality, as evidenced by a substantial increase in gapless and perfectly mapped reads and improved somatic variant calling accuracy. Given that most germline variants of an individual are shared within the human population, the pangenome graph, which represents population-level germline variation, enables more accurate read alignment than GRCh38. To be compatible with linear variant callers, the alignments were converted from the graph to linear (GRCh38) reference through surjection. Surjection partitions an alignment: the portion aligned to reference nodes remains unchanged, while the portion aligned to non-reference nodes is projected onto nearby reference nodes via local alignment.^12^ This process guides reads to correct regions and improves somatic SNV and indel calling accuracy, especially in difficult and extreme regions. We demonstrated that, in some repetitive regions, such as transposable elements, the pangenome becomes important for detecting variants correctly (Figure 3B). For optimal SV detection, after converting graph alignments to linear alignments (surjection), reads need to be indel realigned and polished. However, to date, a tool to perform alignment polishing akin to Abra2^36^ has not been developed for long reads. Furthermore, we observed that the pangenome provides less benefit for accurately aligning long reads compared to short reads, likely because long reads span much larger genomic regions (Figure S6). These factors may collectively explain the observed comparable performance in SV calling between GRCh38 and the pangenome.

Surjecting reads to GRCh38 limits the advantages of the improved representation and completeness of the pangenome graphs. Alternatively, we used the pangenome graph to impute a personalized subgraph containing a pseudo-diploid assembly that closely approximates the DSAs. Calling somatic variants based on the pseudo-diploid assembly yields notable improvements in alignment and in the accuracy of calling somatic SNVs, indels, and SVs, comparable to those achieved with DSAs and exceeding those from the pangenome surjection approach.

Moreover, our study utilized the HapMap mixture, an artificial system that uses germline variants to simulate somatic variants. This has advantages, as the density of germline variants is much higher than that of true somatic variants, enabling benchmarking across the entire genome, including easy, difficult, and extreme regions. The controlled VAF in the mixture allows for benchmarking at various VAFs.

We demonstrated the advantages of pangenome-inferred DSA for reducing germline contamination in somatic variant detection when a matched normal control is unavailable or inapplicable, e.g., normal tissue. In the HapMap mixture, most homozygous HG005 variants were filtered out by pangenome-inferred DSAs. The pangenome enables filtering germline variants at the alignment step, substantially reducing false positives in the initial call set and alleviating the burden on somatic variant callers of distinguishing somatic variants from a large number of germline variants. In contrast, the widely used Panel of Normals (PoNs) approach filters germline variants after variant calling. PoNs are limited in variant type, as they are widely used for SNVs and small indels but not for structural variants^18^. Pangenome includes assemblies from diverse ancestries, and the filtering is not restricted to any variant type. Our study provides proof of principle that using a representative reference can filter out germline variants during the alignment process, effectively serving as an additional layer of germline and somatic variant classification.

We found that the pangenome-inferred DSA enables more comprehensive profiling of somatic variants in a cancer cell line. Using the pangenome, we identified 4,290 (10.15%) novel SNVs in COLO829. Mutational signature analysis showed that these SNVs are highly enriched for the UV light exposure signature, underscoring their biological relevance in melanoma. We showed that genes with low mappability in GRCh38 are well resolved by the pangenome, emphasizing that a more representative reference—such as a pangenome—is essential for confident read alignment and accurate variant detection for these genes. 51.75% of the SNVs unique to the pangenome-inferred DSA were located in regions that are absent or poorly represented in GRCh38. These SNVs arise from insertions, copy−number gains, and other complex genomic loci, including satellite regions. We showed that GRCh38 underestimates the mutation rate in satellite regions. However, even with improved references, mutation rates in these highly repetitive regions are likely still underestimated. Accurate characterization will require further advances in sequencing technologies and alignment algorithms capable of handling highly repetitive genomic regions.

Many newly discovered somatic SNVs using the pangenome are located in melanoma-associated genes such as GNG10^37^, ARHGEF5^31^, and TMEM87b^32^, as well as in genes implicated in other cancer types, such as LINC01410^38^, TRIOBP^33^, FCGBP^34^, and NBPF1^35^. 11 sSNVs were identified as missense mutations, with 2 predicted to be pathogenic by AlphaMissense^30^ (Table S4). 10 of them were located in genes previously reported to be associated with melanoma or other cancer types (Table S4). TMEM87b is the paralog of TMEM87a. TMEM87a was reported as a component of a novel mechanoelectrical transduction pathway and modulates melanoma adhesion and migration^32^. NEB is a prognostic marker in renal cancer based on TCGA^29^. ZSCAN23 is involved in transcription regulation and is a tumor suppressor in pancreatic ductal adenocarcinoma^39^. ARHGEF5 is a key regulator of many cellular events triggered by extracellular signals acting through G protein-coupled receptors and likely contributes to the regulation of cytoskeletal organization^40^. ARHGEF5 promotes epithelial-mesenchymal transition in colorectal cancer^41^, and its overexpression is associated with poor prognosis in acute myeloid leukemia^42^.

LDLRAD4 is involved in the negative regulation of cell migration and epithelial to mesenchymal transition^43^. Decreased LDLRAD4 expression leads to metastasis and poor prognosis in colorectal cancer^44^. ZNF536 is a highly conserved zinc-finger protein that negatively regulates neuronal differentiation by repressing retinoic-acid-induced gene transcription^45^. It has also been linked to poor prognosis in neuroblastoma^46^. Retinoic acid has been reported to affect proliferation in melanoma cell lines^47^. The missense mutation of ZNF536 identified in this study occurs within its C2H2 zinc-finger domain and is predicted to be pathogenic, potentially altering its DNA-binding capability and impacting cell proliferation in COLO829. NBPF1 has been implicated in several types of cancers, including neuroblastoma^35^ and adrenocortical carcinoma^48^. TRIOBP regulates actin cytoskeletal organization, cell spreading, and cell contraction. TRIOBP is highly expressed across cancers, and specific isoform expression is associated with gastric, rectal, pancreatic, and brain cancers^49^. FCGBP is involved in the maintenance of the mucosal structure as a gel-like component of the mucosa and is a prognostic or diagnostic marker for multiple cancer types^50^. LIMS4 lacks literature support of a cancer gene role, likely due to low mappability in GRCh38, and is predicted to be located in the cytoplasm, focal adhesion, and plasma membrane^43^, which could be associated with melanoma metastasis. Collectively, these findings highlight the advantages of using a more representative reference for somatic variant detection and reveal somatic mutations that may be biologically consequential yet missed when relying on GRCh38. This expanded cancer gene mutation catalog may hold new promises in understanding melanoma biology, as well as in developing diagnosis and therapeutics. Pangenome can serve as a great and cost-effective resource for obtaining an individual diploid assembly with reduced reference biases compared with GRCh38.

Together, these results suggest that pangenome improves read mapping and somatic variant calling accuracy, enables efficient germline variant filtering and more comprehensive somatic variant profiling, and opens new avenues to leveraging pangenomic resources for somatic variant detection.

Despite these advances, our study has several limitations. First, the pangenome personalization did not fully capture the germline variants of the true DSAs. More sophisticated genome inference methods need to be developed to further minimize the genetic distance between pangenome-inferred DSAs and true DSAs. Second, current somatic variant callers are based on the linear reference genome, preventing full utilization of pangenome sequences and effective filtering of heterozygous germline variants. Graph-based variant callers are needed to address these limitations. Third, the HapMap mixture uses germline variants to simulate somatic variants. The mutational load and pattern may be different from true somatic variants.

## Methods

### HapMap mixture design and generation

To mimic the somatic variants with a broad spectrum of variant allele fractions (VAFs), six well-studied HapMap samples (HG005, HG02622, HG002, HG02257, HG02486, and HG00438) are chosen to construct the cell mixture samples according to certain ratios. The target composition for the pooled sample was 83.5% HG005, 10% HG02622, 2% each of HG002, HG02257, and HG02486, and 0.5% HG00438, among which HG005 is taken as the germline source, while five additional individuals contribute reads simulating artificial somatic variants.

Each cell line was thawed and seeded into T25 flasks containing 10 mL of culture medium composed of 84% RPMI-1640, 15% FBS, and 1% Glutamax. A portion of the cell suspension was used for counting, and the culture volume was adjusted as needed to maintain a viable cell density between 2×10⁵ and 5×10⁵ cells/mL. Cells were subsequently passed every 3–5 days by dilution based on cell counts until the desired cell number was achieved. Viable cell counts were determined using a Vi-Cell Blu cell counter (Beckman Coulter) with trypan blue staining.

On the day of harvest, cells from each line were pooled into a 250 mL conical tube and mixed thoroughly. Approximately 10 mL of the mixture was transferred to a T25 flask, and an aliquot was taken for cell counting. Sterility was assessed by streaking two sheep blood agar plates, which were incubated at 30 °C and 37 °C for two weeks. Based on the cell counts, the required volume of each cell line was calculated to achieve the target cell number. These volumes were combined in a secondary tube to generate the final pooled suspension. The pooled cells were centrifuged at 228 RCF for 10 minutes, resuspended in 800 mL of cryopreservation medium (65% RPMI-1640, 30% FBS, 5% DMSO), and aliquoted into cryovials at a concentration of 5×10⁶ cells per vial. Cryopreservation was performed using a controlled-rate freezer, and vials were stored in liquid nitrogen vapor.

### HapMap mixture sequencing data generation

The WashU-VAI Genome Characterization Center produced the 500× short-read WGS data using Illumina NovaSeq X Plus and 100× long-read WGS data using PacBio Revio for HapMap mixture cell lines.

### Short-read sequencing

Genomic DNA was quantified using a Qubit Fluorometer. Approximately 600-1,000 ng of DNA was sheared on a Covaris LE220 instrument to generate ∼375 bp fragments. Size selection was performed with a 0.8× ratio of AMPure XP beads (Beckman Coulter) to remove fragments shorter than 300 bp. Dual-indexed libraries were prepared using the KAPA Hyper PCR-free Library Preparation Kit (Roche Diagnostics, Cat #7962371001), incorporating full-length custom adapters (IDT) in a UDI/UMI configuration, featuring 10 bp unique dual indexes (UDIs) and a 9 bp unique molecular identifier (UMI) in the i7 position. Library molarity was quantified using the KAPA Library Quantification Kit (Roche Diagnostics). Libraries of 150 bp paired-end reads were sequenced on the NovaSeq X platform. To achieve >500× coverage, four independent libraries were prepared and sequenced, and the resulting data were pooled from 4 library sources.

Library molarity was accurately quantified by qPCR using the KAPA Library Quantification Kit (KAPA Biosystems/Roche) following the manufacturer’s instructions, ensuring optimal cluster density for sequencing on the Illumina NovaSeq X Plus platform. Normalized libraries were then sequenced on a NovaSeq X Plus flow cell using a 151×10×10×151 paired-end sequencing protocol, generating whole-genome sequencing coverage exceeding 500×.

### Long-read sequencing

PacBio HiFi SMRTbell libraries were prepared based on the Procedure & Checklist – Preparing Whole Genome and Metagenome Libraries Using SMRTbell Prep Kit 3.0 (PacBio). Genomic DNA was initially treated with DNAFluid+ (P/N E07020001) at a speed setting of 40 on the Diagenode Megaruptor 3 to dissociate aggregates and homogenize the sample. The homogenized DNA was then sheared twice using the Shearing Kit (P/N E07010003) at speeds of 28 and 30, targeting a fragment size mode of ∼20 kb. DNA quality and fragment size distribution were assessed using the Qubit High Sensitivity DNA Kit and the Agilent Femto Pulse system (Genomic DNA 165 kb Kit).

Library preparation followed the PacBio protocol and included barcoded adapters from the SMRTbell Barcoded Adapter Plate 3.0 (PacBio, P/N 102-009-200) for multiplexed sequencing. Libraries were size-selected on the Sage PippinHT using the 0.75% Agarose High-Pass 75E Kit (P/N HPE7510) with a lower cutoff between 15,000–17,000 bp. Final library preparation for sequencing was performed according to instructions generated by SMRT Link v13.0, using the PacBio Revio Polymerase Kit (P/N 102-817-600).

Sequencing was conducted on the PacBio Revio system using 30-hour movie collections with an on-plate concentration of 170–200 pM. To achieve ∼100x coverage (approximately 300 Gb total), three SMRT Cells were used, with average Q-scores of Q33, Q33, and Q32, respectively.

### COLO829 and COLO829BL sequencing data and COLO829BL DSA generation

The PacBio HiFi sequencing data of COLO829 (SMAFIJPL47BQ, SMAFIO69TOMB, SMAFIURDW1P6) and COLO829BL (SMAFIFRLAQ54) cell lines were generated as part of the SMaHT benchmarking study^51^. COLO829BL DSA was generated by UWSC GCC^24^ using Verkko2^23^. All data were downloaded from the SMaHT data portal (https://data.smaht.org).

### Pangenome graph construction and characterization

The assemblies from HPRC release 1^12^, GRCh38, and CHM13 were constructed into v2 and v3 pangenome graphs using Minigraph-Cactus (v2.9.7)^15^, with GRCh38 as reference. Nodes, edges, and edge degrees of three pangenome graphs were counted using Panacus (v0.2.4)^52^. The clipped and allele-filtered graphs were used for alignment and surjection. The clipped and unfiltered graphs were used for pangenome personalization. Minigraph-Cactus also generated the pangenome deconstructed VCF, which was used to visualize non-reference variants in Figure 3B.

### Personalized pangenome (pangenome-inferred DSA) construction and evaluation

We used the HG005 simulated short-read WGS data and COLO829BL short-read WGS data to construct the personalized pangenome graphs from v3 pangenome graphs (clipped and not allele-filtered). We indexed the clipped graph using vg (v1.64.1) to create the index .hapl file, .ri file and .dist file:

vg index -t 10 -j $dist --no-nested-distance $gbz
vg gbwt -p --num-threads 10 -r $ri -Z $gbz
vg haplotypes -v 2 -t 10 -H $hapl $gbz

and utilized the KMC (v3.2.4)^53^ with -k29 -m128 -okff to count the k-mers from the sample’s WGS library. vg haplotype^17^ was used for haplotype sampling and constructing the personalized pangenome graph.

vg haplotypes -v $level -t $threads --include-reference --diploid-sampling -i $hapl -k $kff -g $output_gbz $gbz

This imputes a subgraph from the pangenome graphs with GRCh38 as reference and two recombinant assemblies (hap1, hap2). Two recombinant assemblies were extracted from graphs to FASTA format and used as pangenome-inferred assemblies.

The pangenome-inferred assemblies are evaluated using linear and graph-based approaches. The linear approach compares the number of base substitutions, insertions, and deletions between pangenome-inferred assemblies with HG005 maternal/paternal assemblies and that between GRCh38 and HG005 maternal/paternal assemblies. The assemblies were aligned with Minimap2 (v2.28)^20^ using -cx asm5; --secondary=no settings to generate PAF files. The paftools.js stat was used to count the number of substitutions, insertions, and deletions from PAF files. The graph-based approach incorporated pangenome-inferred assemblies and the HG005 or COLO829BL diploid assemblies^24^ into a five-assembly Minigraph-Cactus^15^ validation graph, with GRCh38 as reference. The graphs were deconstructed to VCF with genotypes from each assembly, which were then used to compare genotypes between the pangenome-inferred assemblies and the HG005 or COLO829BL diploid assemblies and calculate the capture rate.

### Assembly scaffolding

RagTag (v2.1.0)^54^ was used to scaffold the assemblies (HG005 maternal and paternal assemblies, pangenome-inferred assemblies of HG005 and COLO829BL), using GRCh38 (GCA_000001405.29) as the reference assembly.

### Whole-genome sequencing data processing for the HapMap mixture and COLO829

After raw sequence quality assessment, Illumina short reads were processed using fastp^55^ to remove read pairs containing polyG artifacts. Illumina short reads of the HapMap mixture were aligned to linear references: GRCh38, HG005 paternal or maternal assemblies, and graph-based references: v1, v2, and v3 pangenome graphs (clipped and allele-filtered) and a personalized pangenome graph (subgraph contains GRCh38, pangenome-inferred assemblies of HG005 hap1 and hap2). GRCh38 (GCA_000001405.15) excludes all the ALT contigs and Human decoy sequences from version GCA_000786075.2 of GRCh38, including chromosomes from the GRCh38 primary assembly unit, mitochondrial genome from the GRCh38 non-nuclear assembly unit, unlocalized scaffolds from the GRCh38 primary assembly unit, and unplaced scaffolds from the GRCh38 primary assembly unit and Epstein-Barr virus (EBV) sequence. HG005 paternal and maternal assemblies were obtained from HPRC release 1^12^. chrX and chrM from HG005 maternal assembly were added to HG005 paternal assembly, and chrY from HG005 paternal assembly was added to HG005 maternal assembly to ensure the assemblies have a full set of chromosomes. The PacBio HiFi data of the HapMap mixture were aligned to linear references: GRCh38, HG005 paternal or maternal assemblies, and pangenome-inferred assemblies of HG005 hap1 or hap2. The PacBio HiFi data of COLO829 and COLO829BL were aligned to linear references: GRCh38, pangenome-inferred assemblies of COLO829BL hap1 or hap2.

Illumina short read data were aligned to the linear references using BWA-MEM (v0.7.17)^56^. PacBio HiFi data were aligned using pbmm2^20^ (v1.13.1). The short-read data were aligned to the genome graphs using vg (v1.64.1) giraffe^16^. The reads were then surjected to GRCh38 (for v1, v2 and v3 pangenome graphs) or pangenome-inferred assemblies of HG005 (for personalized pangenome graph) using vg surject^12^. For short-read data, the alignments were processed with Mark Duplicate, Indel Realignment, and Base Quality Score Recalibration (BQSR) using GATK (v4.1.0.0)^57^ before variant calling.

### Somatic variant calling and filtering for the HapMap mixture

SNVs and indels (small insertions and deletions less than 50 base pairs) were called from 500x short-read data using MuTect2 (v4.5.0.0)^18^ tumor-only mode. The variants are then filtered using MuTect2 FilterMutectCalls^18^. The “haplotype” or “clustered_events” were accepted in addition to PASS FILTER tags to match the characteristics of the HapMap mixture, which has more than 10 million SNVs from 6 different cell lines and more than 6 million in the somatic VAF range, much more than expected in a healthy human sample. The SNVs and indels were decomposed and normalized by bcftools (v1.20-24-g5d48dbc3)^58^ norm -a --atom-overlaps ’.’ --check-ref w -N and separated into SNVs and indels by Picard (v2.9.0) SplitVcfs.

SVs (insertions and deletions >= 50 base pairs) were called and filtered from 100x long-read data using Sniffles2 (v2.6.3) mosaic mode and set –mosaic-af-max 0 and –mosaic-af-max 0.218 following the developer’s suggestion.^19^ In Sniffles2 mosaic mode, the ALN_NM filter removes artifacts by excluding SVs whose reads show an unusually high length-weighted mismatch rate relative to the dataset average.^19^ This filter was disabled because reduced genome-wide alignment mismatch rates in DSAs or pangenome-inferred DSAs (due to decreased reference bias) increase the filter stringency, making it incomparable with GRCh38.

### Alignment metrics

The mapped reads, properly paired reads, gapless reads, and perfectly mapped reads were used as alignment metrics. Gapless reads are defined as alignments without insertions and deletions and might contain mismatches and/or softclips. Perfectly mapped reads are defined as alignments that map exactly to the reference. Properly paired reads are defined as alignments that paired-end reads must map, be in a faced orientation, and within a reasonable distance.

For alignments to linear references, the short-read alignment metrics were generated using samtools (v1.18)^58^ stats, flagstats, and MD, CIGAR fields in BAM files. The long-read alignment metrics were generated using Qualimap (v2.2.1)^59^. For alignments to genome graphs, the short and long read alignment metrics were generated using vg stats from the GAM file.

### Benchmarking variant sets generation

The benchmarking variant sets of the HapMap mixture based on GRCh38, pangenome-inferred assembly of HG005 (hap1 and hap2), and HG005 maternal and paternal assemblies were generated and validated using a genome-graph approach^14^ (Figure S7). The assemblies of six HapMap samples were obtained from HPRC release 1^12^. The benchmarking variant sets contain all possible SNVs, indels, SVs from five somatic samples in the HapMap mixture, and benchmarking confident regions (assembly-reliable regions defined by HPRC).^14^ The benchmarking confident regions for the pangenome-inferred assemblies of HG005 (hap1 and hap2) and the HG005 maternal and paternal assemblies were lifted over from the GRCh38 confident regions.

### Region difficulty stratification

The GRCh38 stratification into easy, difficult, and extreme regions was generated by the SMaHT network (https://data.smaht.org). The easy regions are based on 1KG strict mask^60^. The difficult regions are regions outside of easy regions and within sample-agnostic easy regions defined using a pan-genome approach^61^. The extreme regions are defined as those outside the easy and difficult regions. The stratifications for pangenome-inferred DSA of HG005, HG005 maternal and paternal assemblies were generated by lifting over regions from GRCh38 to each assembly using UCSC liftOver with default settings. The CHAIN files were generated by the nf-LO pipeline (v1.8.6)^62^ with default settings. These regions are then overlapped with confident regions from benchmarking variant sets using bedtools^63^ -isec.

### Variant calling benchmark

The benchmarking was performed within the benchmarking confident regions of chr1-22, chrX, and chrY. SNVs and indels were benchmarked using RTG (v3.12.1) vcfeval^64^ with –squash-ploidy and –all-records. SVs were benchmarked using Truvari (v5.2.0)^65^ with --pctsize 0.9, --pctseq 0.9, --pick multi.

The somatic SNVs, indels, and SVs were partitioned into 9 VAF bins: <0.5%, 0.5-1%, 1-1.5%, 1.5-2%, 2-3%, 3-4%, 4-5%, 5-10%, 10-16.5%. For recall, the benchmarking variant set was split by VAF, and each bin was compared with the entire call set, while for precision, the call set was split by VAF and compared with the entire benchmarking variant set. The precision and recall were calculated as: Precision = TP/(TP+FN); Recall = TP/(TP+FP).

### Somatic variant calling and mutational signature analysis for COLO829

Somatic SNVs were identified using DeepSomatic (v1.8.0)^22^ tumor–normal mode from COLO829 and COLO829BL PacBio HiFi data. The somatic SNV calling was performed using GRCh38, pangenome-inferred DSA hap1 or hap2 of COLO829BL as the reference. The SNV calls with more than one alternative allele supporting reads in COLO829BL BAM were removed in the initial call sets. Next, the SNV calls from hap2 were lifted over to hap1 and compared with the hap1 call set to remove the duplicated SNV calls. This harmonized call set was lifted over to GRCh38 and compared with GRCh38 SNV calls. The lifted over SNVs that did not overlap with GRCh38 calls and SNVs that failed to lift over were defined as pangenome-inferred DSA unique SNV calls. Mutational signature analysis was performed on unique and shared SNV calls from GRCh38 or pangenome-inferred DSAs with SigProfilerMatrixGenerator (v1.3.3)^66^ and SigProfilerAssignment (0.2.3)^67^ to obtain SBS and DBS profiles and signatures based on COSMIC (v3.4)^68^ (https://github.com/ryansohny/VCF2SPECTRUM).

### Somatic SNV annotation and pathogenicity prediction for COLO829

The somatic SNVs on GRCh38 were annotated with VEP (v115.2)^70^. The AlphaMissense^30^ pathogenicity scores were generated using VEP plugins. To annotate SNVs that cannot be lifted over from pangenome-inferred DSA to GRCh38, the GRCh38 GENCODE annotation (v49)^71^ was mapped to pangenome-inferred DSA with LiftOff^72^ with the following command:

liftoff -p 16 -g gencode.v49.primary_assembly.basic.annotation.gtf -o hs_hprc-v1.1-mc-chm13_20x_1_gencode.v49.primary_assembly.basic.annotation_copies.gtf -u
unmapped_features.hs_hprc-v1.1-mc-chm13_20x_1_.txt -cds -copies hs_hprc-v1.1-mc-chm13_20x_1.fasta ../GRCh38.primary_assembly.genome.fa

The somatic SNVs were then annotated with BCFtools/csq^73^ to obtain variant consequences (missense/synonymous). The k100 mappability for GRCh38 and pangenome-inferred DSA was generated by GenMap^74^ with the following commands:

genmap index -F hs_hprc-v1.1-mc-chm13_20x_1.fasta -I hs_hprc-v1.1-mc-chm13_20x_1.genmap
genmap map -K 100 -E 0 -I hs_hprc-v1.1-mc-chm13_20x_1.genmap -O genmap_out --wig

### SNV mutation rate analysis for COLO829

Pangenome-inferred DSA of COLO829 was annotated using RepeatMasker (v4.1.9)^69^ using default settings. The RepeatMasker annotation of GRCh38 was obtained from https://www.repeatmasker.org/species/hg.html. The somatic SNVs in repetitive elements were counted based on RepeatMasker annotations. The somatic SNVs in the exon or intron were counted based on VEP annotations. The SNV mutation rate was calculated by dividing the total SNV counts by the total region length.

## Supporting information

All_supplimentary_figures_tables

## Data and code availability

The whole genome sequencing data of HapMap mixture, COLO829, and COLO829BL, and COLO829BL DSA generated by the SMaHT network can be found at https://data.smaht.org/. The HapMap mixture benchmarking variant sets based on GRCh38, pangenome-inferred assembly of HG005 (hap1 and hap2), and HG005 maternal and paternal assemblies can be found in the repository on Zenodo: https://zenodo.org/records/17544617. All scripts are available at: https://github.com/twlab/SMaHT_Pangenome_Benchmark.

## Acknowledgements

We thank Genome Technology Access Center at McDonnell Genome Institute at Washington University School of Medicine for sequencing data generation and management. We also thank all members in the Somatic Mosaicism across Human Tissues Network. These studies were supported through NIH grants UM1DA058219, 3UM1DA058219-01S1, U24NS132103, UM1HG010971, and U41HG010972.

## Author contributions

Q.F., Z.X., J.F.M., and T.W. contributed to study design and conceptualization. R.F., B.K.J., M.F.M., T.P., and H.J.J. performed cell culture and sequencing data production. T.L. managed sequencing data producing pipeline. C.T. and D.L. managed sequencing data transfer. Q.F., Z.X., W.Z., N.K., Z.T., A.R., J.F.M., B.M., D.A., E.A.B., J.E.G., E.M., J.S., X.Z., S.D., Y.C., Z.L. performed bioinformatics analyses. Q.F. and B.M. generated figures. Q.F., J.F.M., and Z.X. wrote the initial manuscript. T.W., J.F.M., B.M., S.C.J., N.K., and Z.T. reviewed and edited the manuscript. T.W., R.F., and H.S. administered the project. T.W., S.C.J., B.F., H.S., and H.A.L. provided funding and resources.

