## Supplementary material for "Leveraging Human Pangenome for Improved Somatic Variant Detection": All_supplimentary_figures_tables

### Supplementary figures

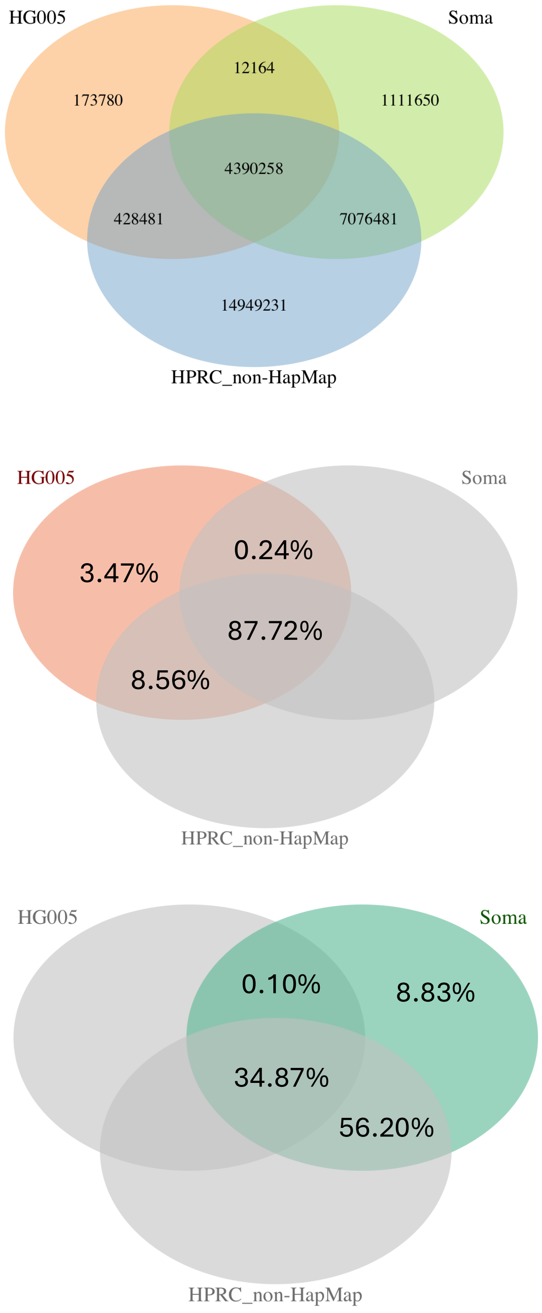


**Figure S1. Characterization of the variants of HG005 and 5 somatic samples shared by other individuals in HPRC release 1 assemblies.** A pangenome graph that contains the assemblies of all 6 HapMap samples, along with other individuals' assemblies in HPRC release 1, was constructed and decomposed to VCF using Minigraph-Cactus. Variants shared by HG005, 5 somatic samples or non-HapMap samples were counted. Most variants (>90%) of HG005 or 5 somatic samples were shared by other individuals. HG005: variants from HG005; Soma: variants from 5 somatic samples in the HapMap mixture; HPRC_non-HapMap: variants from HPRC release 1, except 6 HapMap individuals.

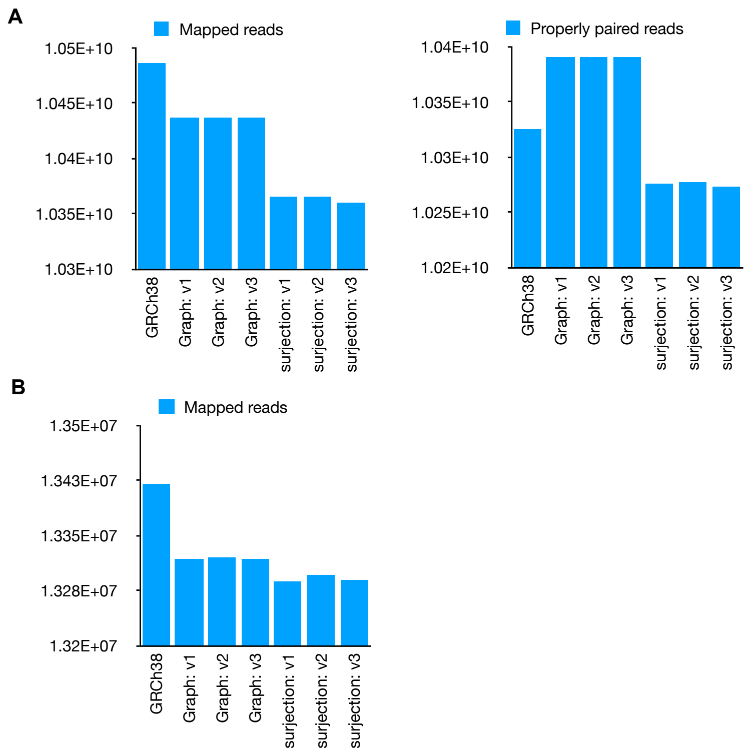


**Figure S2. The number of mapped reads and properly paired reads for HapMap mixture** (**A**) short reads and (**B**) long reads. The pangenome aligns a slightly lower number of reads compared with GRCh38.

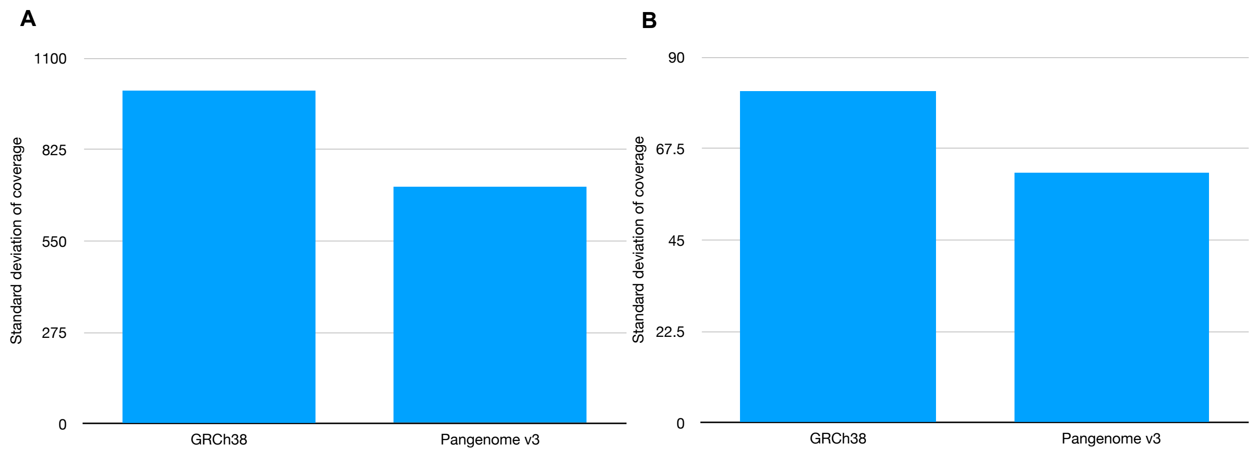


Figure S3. Standard deviation of coverage for GRCh38 and pangenome surjection approach for (**A**) short reads and (**B**) long reads. Standard deviation was calculated from coverage in each 1 kb bin.

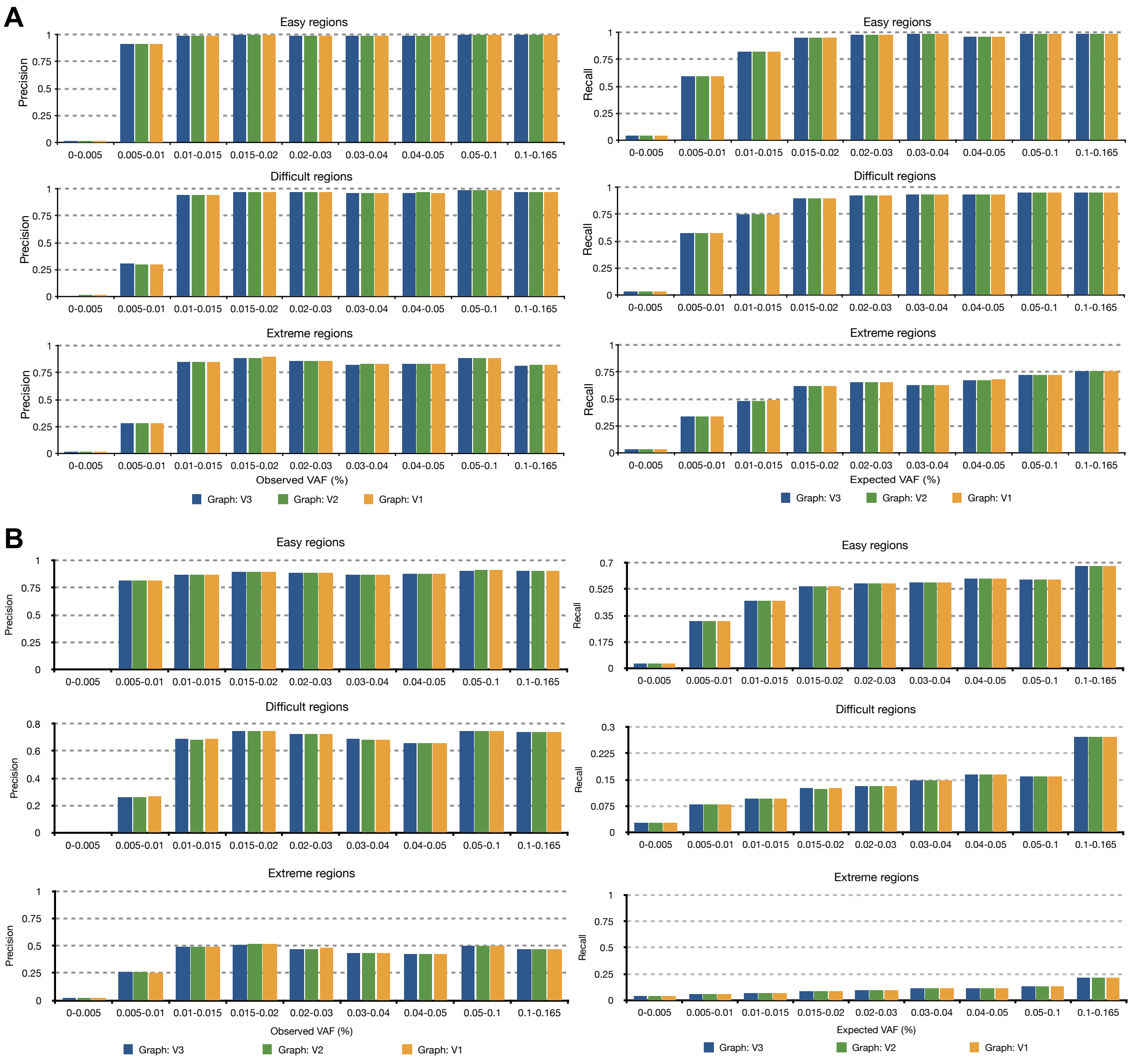


**Figure S4. Comparison of the precision and recall among v1, v2, and v3 pangenome graphs** for (**A**) SNV and (**B**) indel. SNVs and Indels were called from HapMap mixture short reads.

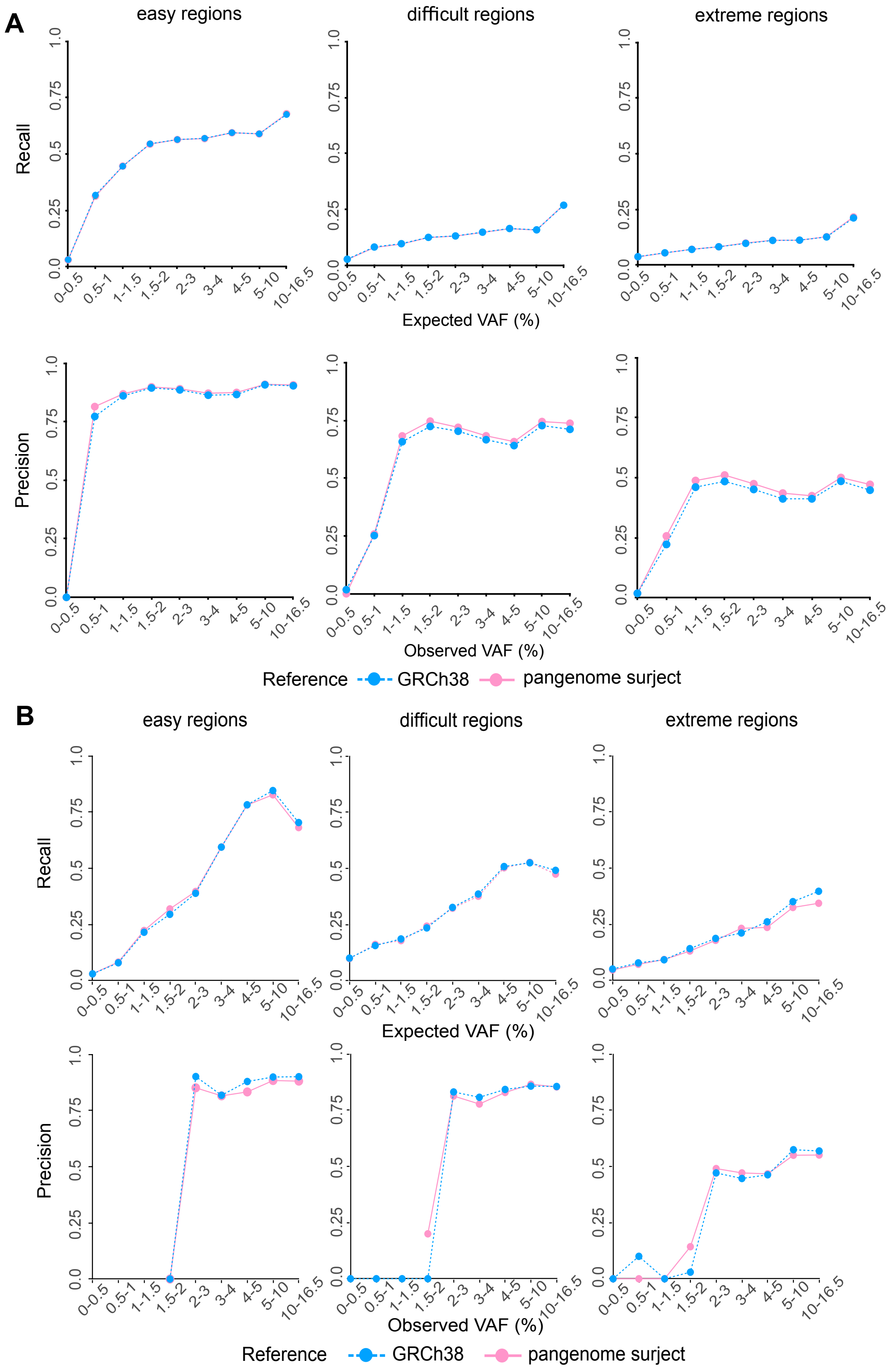

A
easy regions
difficult regions
extreme regions
Recall
Expected VAF (%)
Precision
Observed VAF (%)
0-0.5
0.5-1
1-1.5
1.5-2
2-3
3-4
4-5
5-10
10-16.5
0.0
0.25
0.5
0.75
1.0
Reference
GRCh38
pangenome surject
B
easy regions
difficult regions
extreme regions
Recall
Expected VAF (%)
Precision
Observed VAF (%)
Reference
GRCh38
pangenome surject

**Figure S5. Indels and SVs calling performance for GRCh38 and pangenome surjection approach in the HapMap mixture. (A)** Indels and **(B)** SVs were called from HapMap mixture short and long reads, respectively.

A

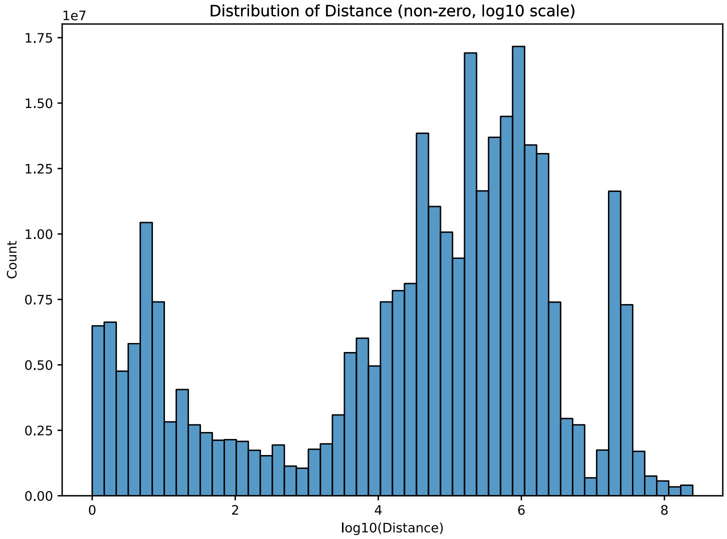


B

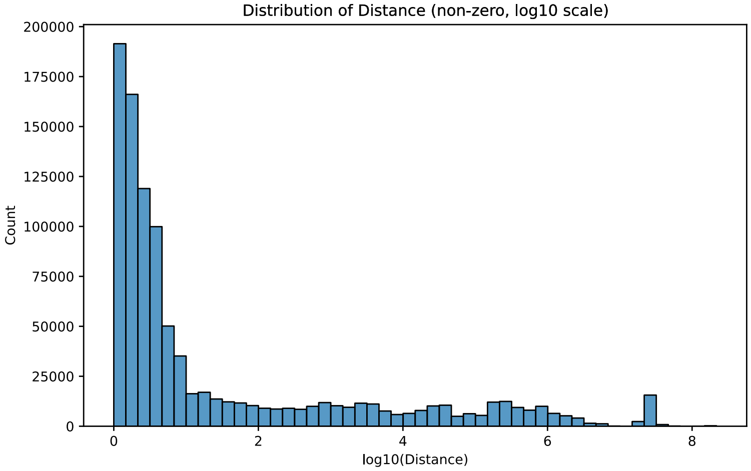


C

| | Illumina short read | PacBio HiFi long read |
|---|---|---|
| Total mapped reads (GRCh38) | 10,486,102,554 | 13,419,614 |
| Reads mapped to different chromosomes | 304,405,196 (2.90%) | 180,394 (1.47%) |
| Reads with distance > 1000 | 229,388,312 (2.19%) | 197,134 (1.47%) |
| Reads with distance > 10,000 | 207,206,369 (1.98%) | 141,226 (1.05%) |

**Figure S6. Distance between GRCh38 alignments and pangenome surjected alignments for Illumina short reads and PacBio HiFi long reads. A**. Distribution of non-zero distances for short reads. **B**. Distribution of non-zero distances for long reads. **C**. Statistics for reads mapped to different chromosomes, reads with distance > 1000 or distance > 10,000 for short reads and long reads. A smaller proportion of long reads are mapped to different chromosomes or distant locations between the two approaches than that of short reads.

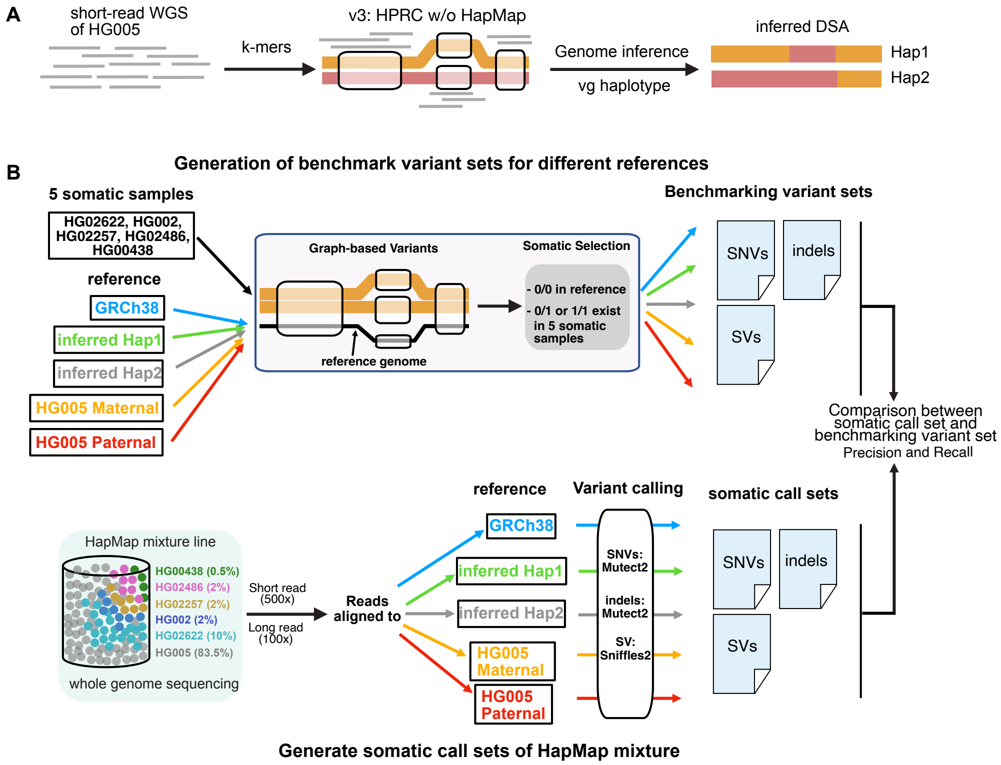


**Figure S7. Schematic workflow of pangenome-inferred DSA construction and its benchmarking. A**. The k-mer counts from short-read WGS of HG005 were used to haplotype sample pseudo-diploid assemblies that approximate DSA (HG005 assembly) from the full, clipped v3 pangenome graph. **B**. The SNV, indel, and SV call sets based on GRCh38, pangenome-inferred DSA hap1 and hap2, and HG005 paternal and maternal assembly were generated and compared with respective benchmarking variant sets.

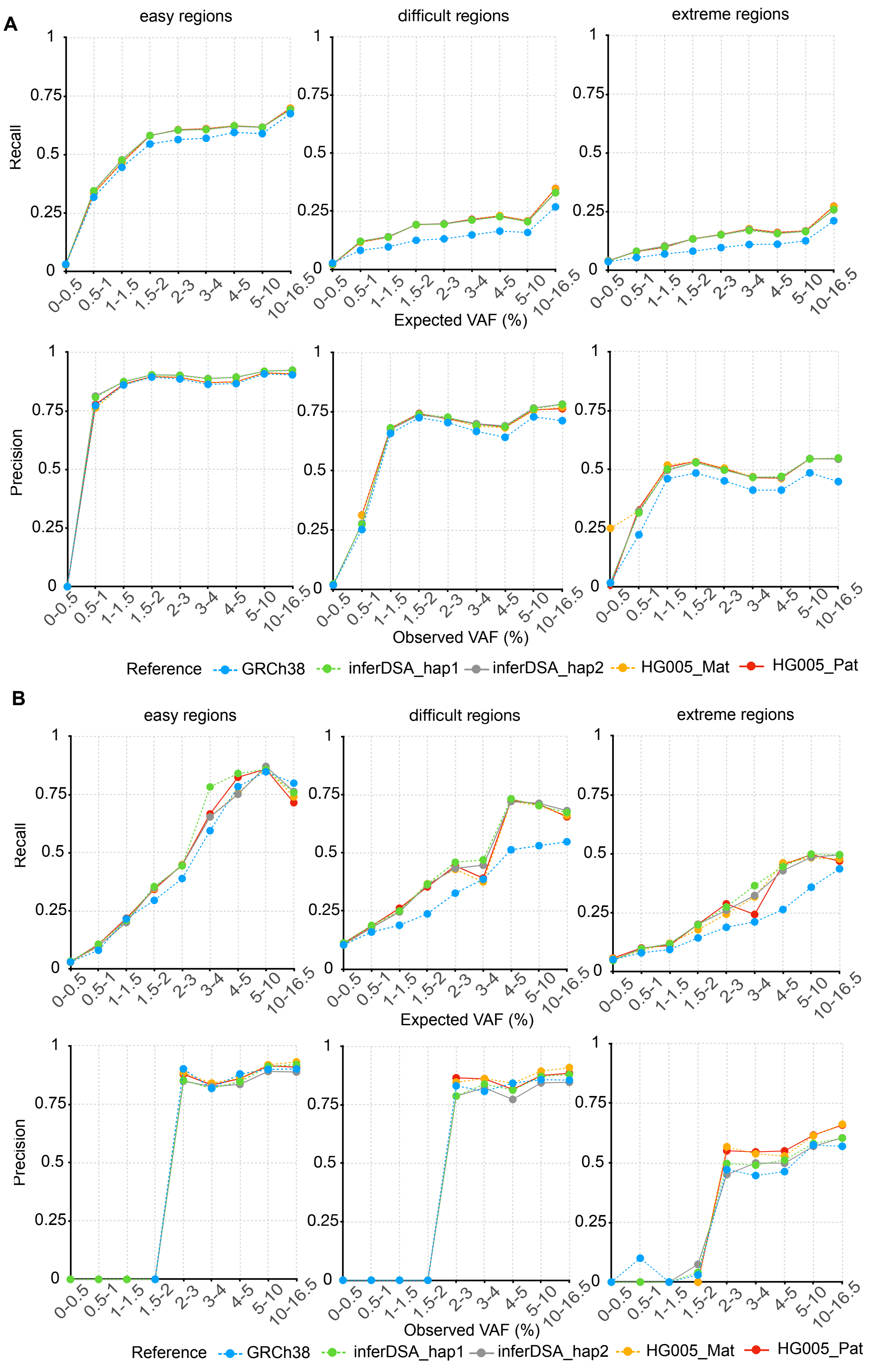
A
easy regions
difficult regions
extreme regions
Recall
Expected VAF (%)
Precision
Observed VAF (%)
0-0.5
0.5-1
1-1.5
1.5-2
2-3
3-4
4-5
5-10
10-16.5
Reference
GRCh38
inferDSA_hap1
inferDSA_hap2
HG005_Mat
HG005_Pat
B
easy regions
difficult regions
extreme regions
Recall
Expected VAF (%)
Precision
Observed VAF (%)
Reference
GRCh38
inferDSA_hap1
inferDSA_hap2
HG005_Mat
HG005_Pat

**Figure S8. Indels and SVs calling performance for GRCh38, pangenome-inferred DSAs (hap1 and hap2), and DSAs (HG005 paternal and maternal assemblies). (A)** Indels and **(B)** SVs were called from HapMap mixture short and long reads, respectively.

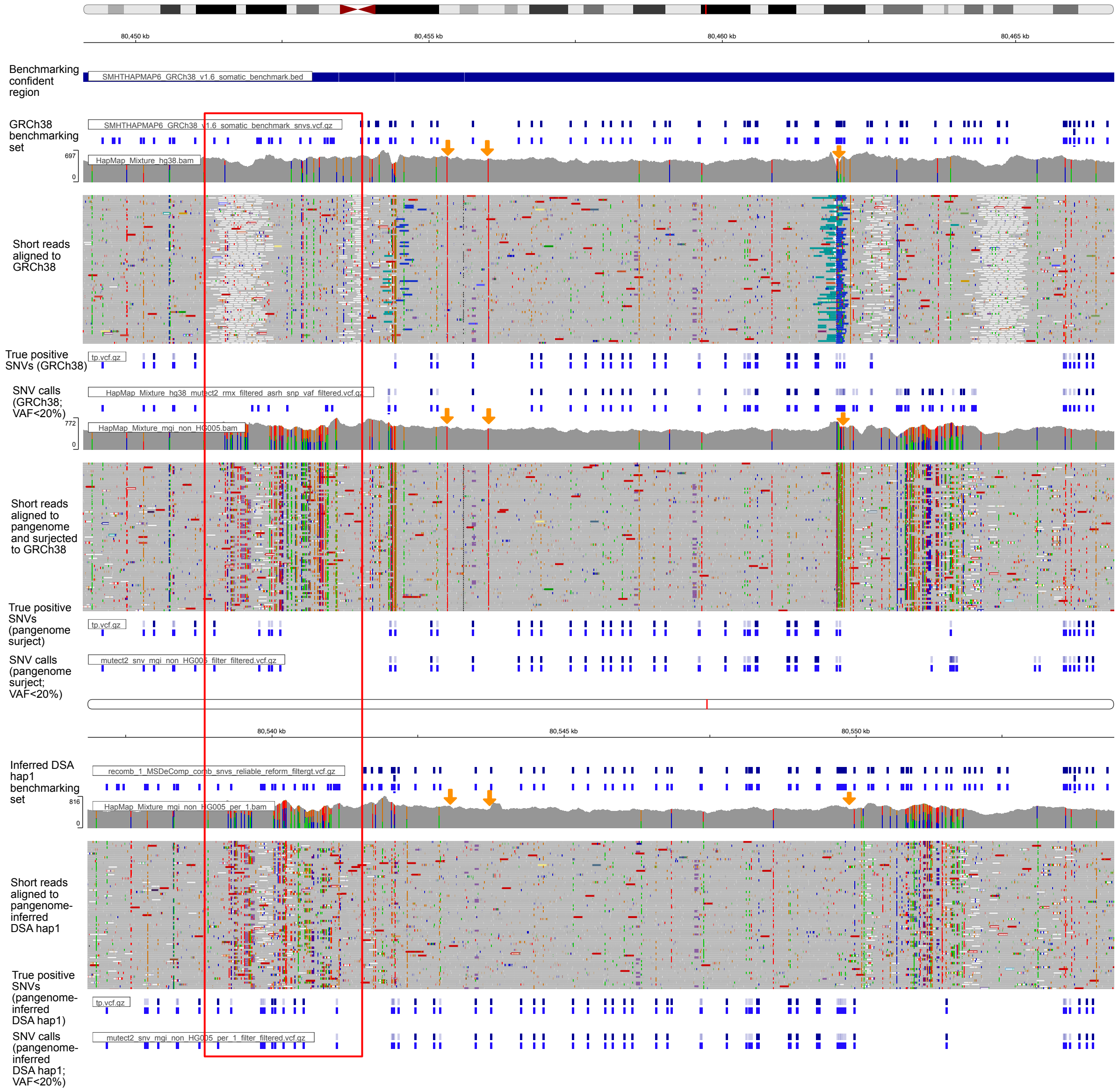


**Figure S9. Example locus (chr12:80,449,114-80,466,689) indicates improved alignment and somatic variant detection on pangenome-inferred DSA.** HapMap mixture short reads were aligned to GRCh38, pangenome (followed by surjection to GRCh38), and pangenome-inferred DSA hap1. Red box: Pangenome-inferred DSA hap1 enables accurate SNV detection in the L1PA3 element. Yellow arrow: homozygous germline variant filtered by pangenome-inferred DSA at the alignment step.

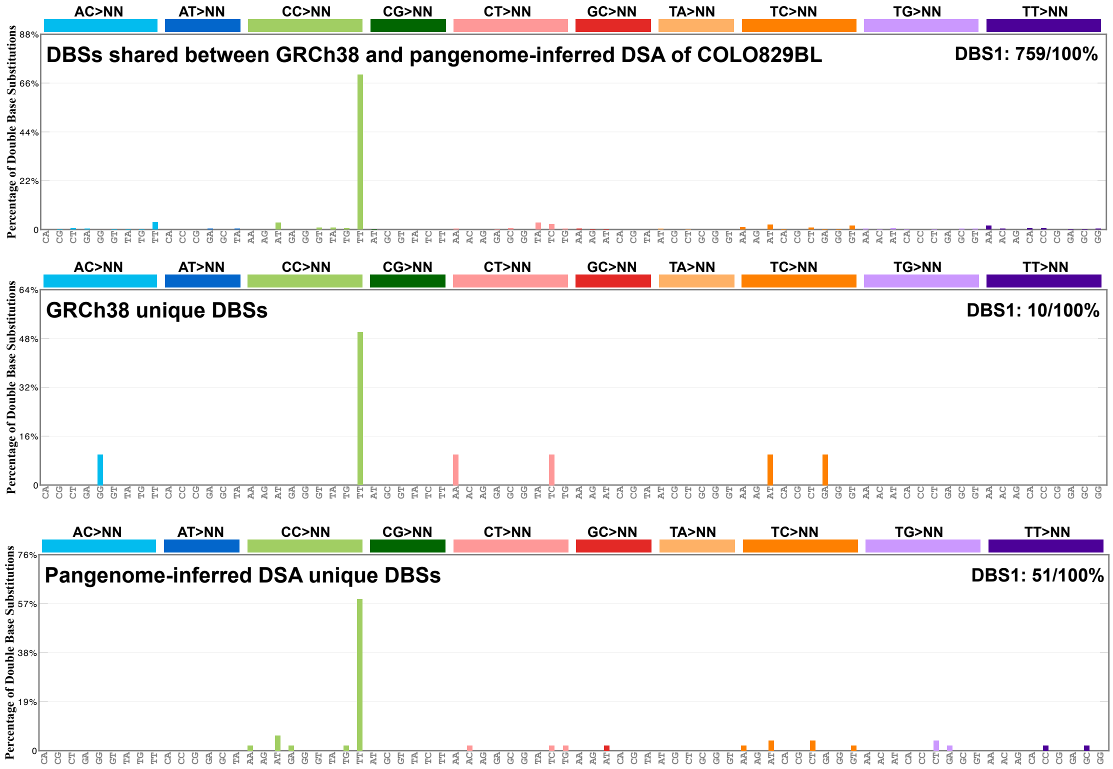


**Figure S10. DBS signatures of shared and unique somatic SNVs from GRCh38 or pangenome-inferred DSA of COLO829BL.** Both shared and unique somatic SNVs were enriched in the DBS1 signature. The pangenome-inferred DSA identifies four times more unique DBS contributing to the DBS1 signature than GRCh38.

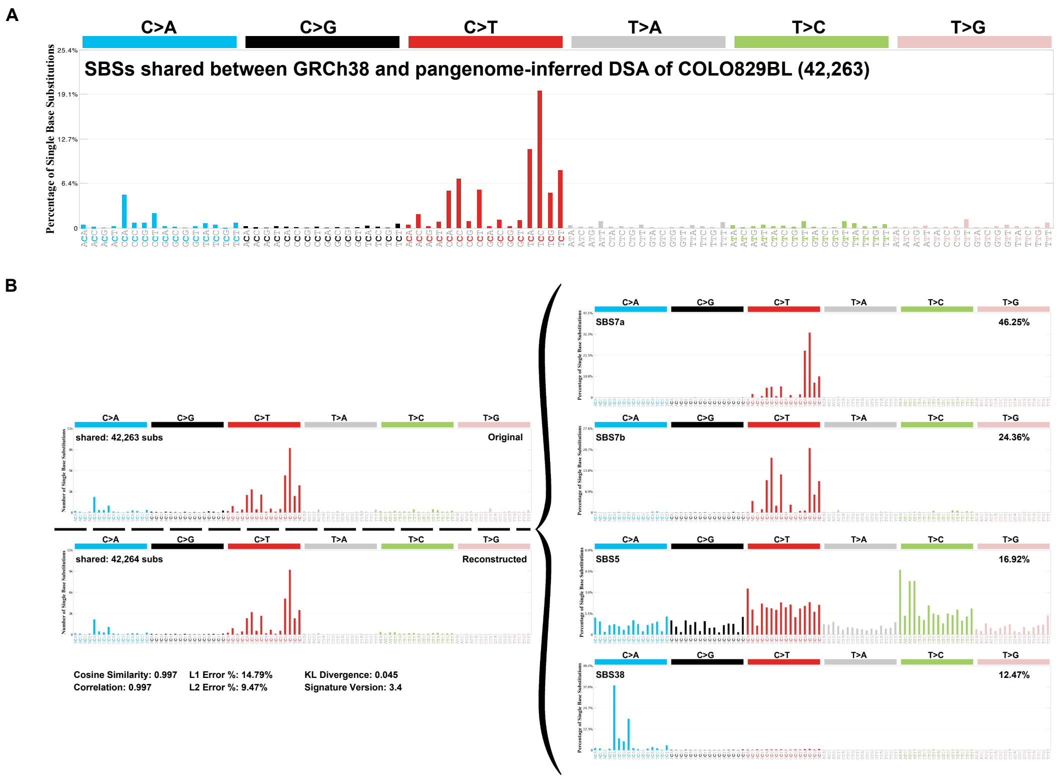


**Figure S11. SBS signatures for SNVs shared between GRCh38 and pangenome-inferred DSA of COLO829BL. A.** somatic SNV mutational profile. **B.** somatic SNV mutational signature.

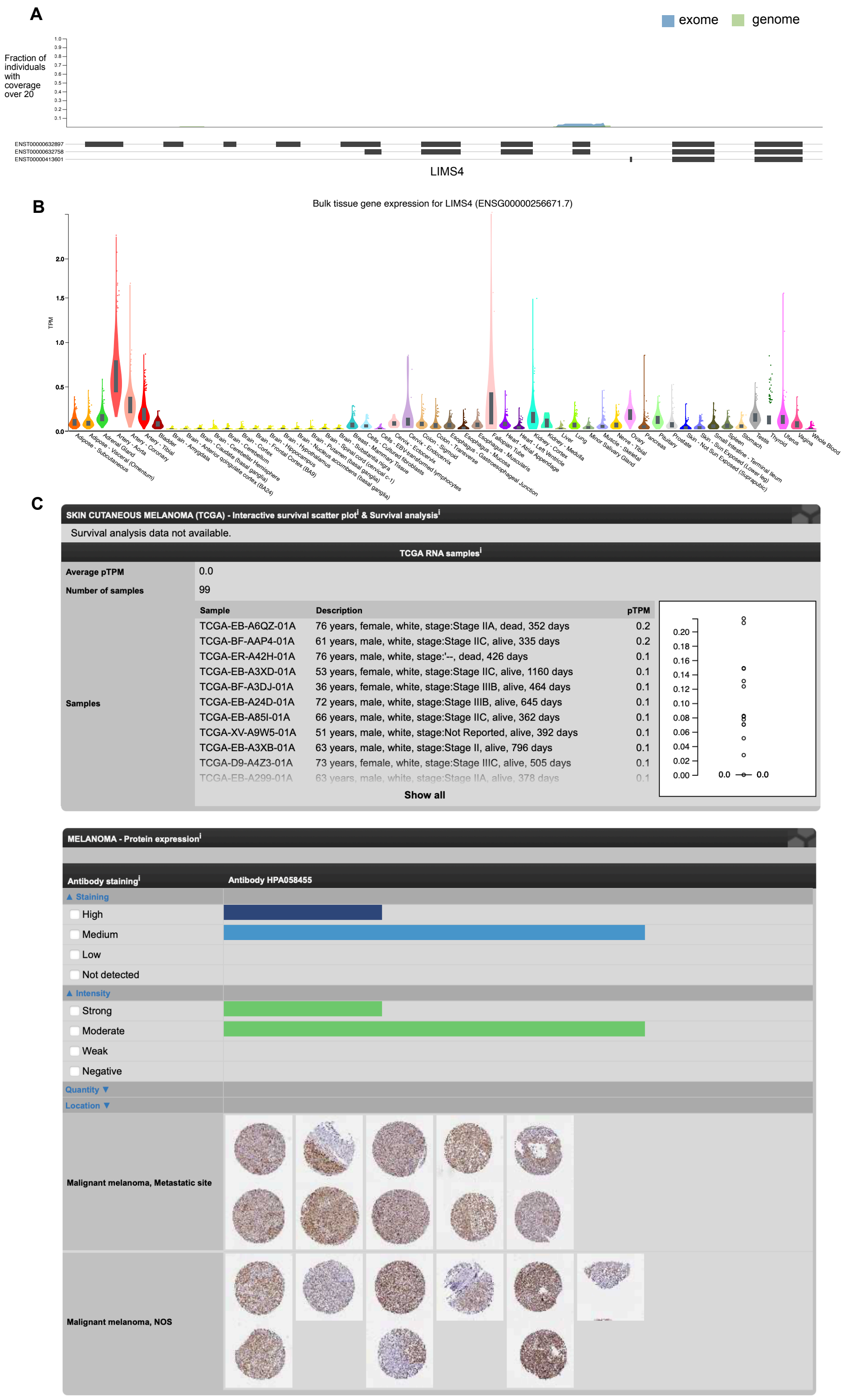


**Figure S12. Population coverage of LIMS4 and its expression across human tissues and melanoma based on open-access databases. A**. LIMS4 population coverage based on gnomAD[1]; **B**. RNA level across tissue based on GTEx[2]; **C**. RNA and protein levels in melanoma based on TCGA[6].

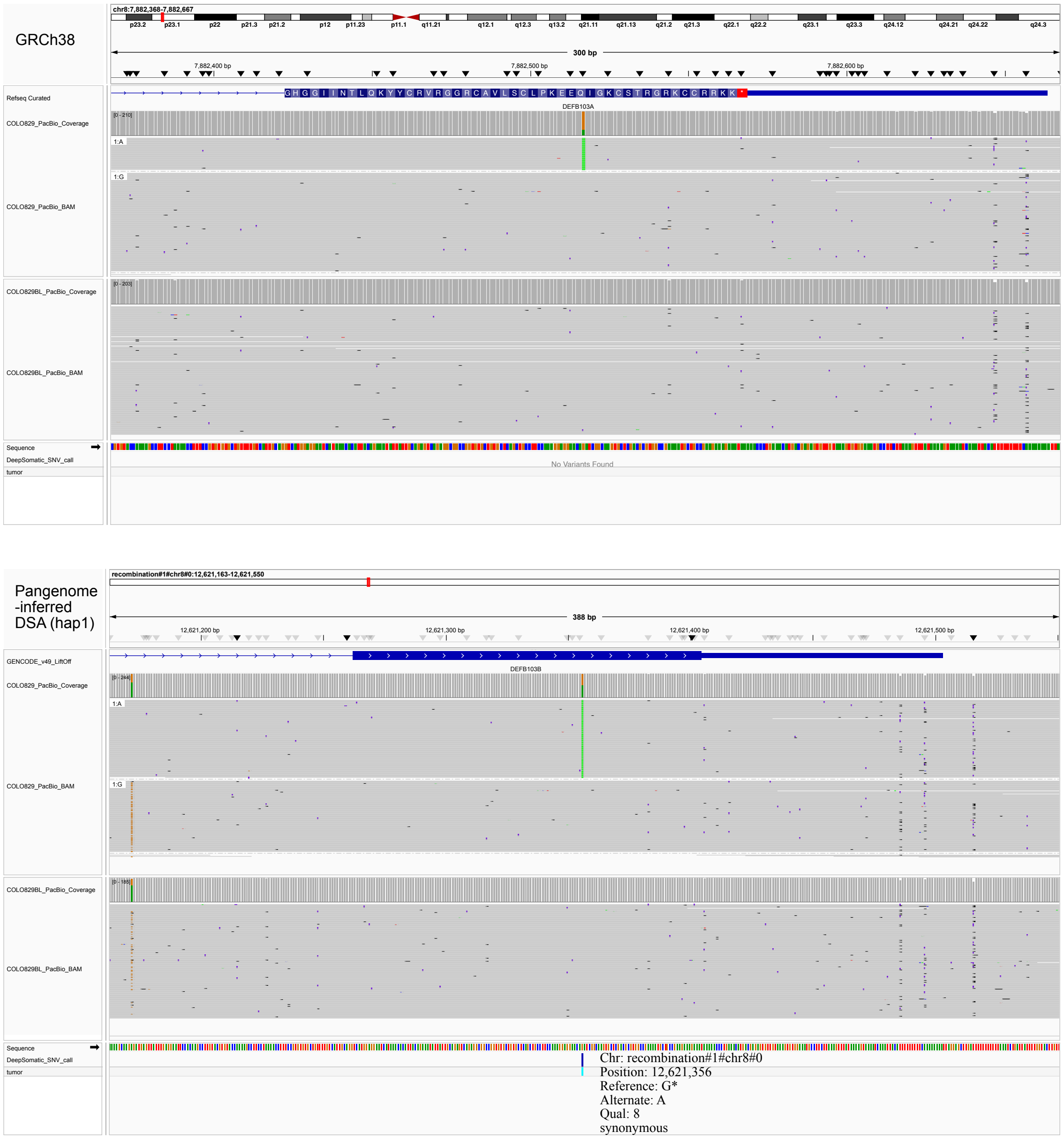


**Figure S13. Examples of somatic SNVs detected by the pangenome-inferred DSA in COLO829BL but missed in GRCh38 and could be lifted over.** The pangenome-inferred DSA aligned a greater number of somatic variant supporting reads, thereby enabling more sensitive and accurate somatic mutation detection in DEFB103B.

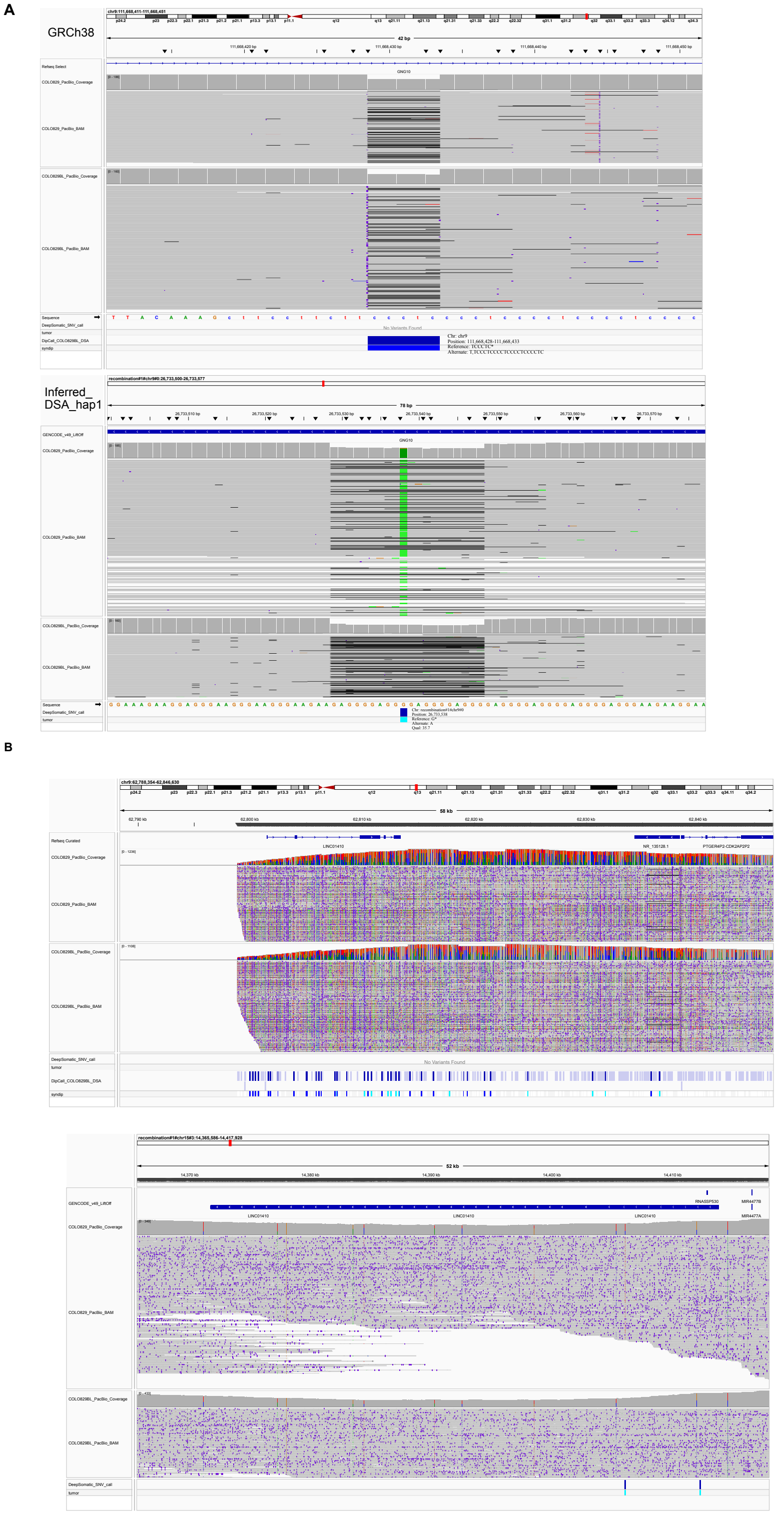


**Figure S14. Examples of somatic SNVs detected by the pangenome-inferred DSA in COLO829BL but missed in GRCh38 and unable to be lifted over. A.** The somatic SNV resides in a simple repeat expansion in the intron of the GNG10 gene in the COLO829BL haplotype. **B.** The copy number gain of LINC01410 in COLO829BL resolved by pangenome-inferred DSA results in reasonable coverage and somatic variant detection.

**Supplementary tables**

| Reference | Covered Reliable Region | | Counts | | | |
|---|---|---|---|---|---|---|
| | Base (Gb) | % genome | SNV | Indel | SV | Total |
| GRCh38 | 2.75 G | 89.1% | 6,037,703 | 1,832,989 | 51,006 | 7,921,698 |
| HG005 maternal | 2.78 G | 92.0% | 7,689,340 | 2,080,542 | 44,546 | 9,814,428 |
| HG005 paternal | 2.78 G | 91.7% | 7,513,048 | 2,062,348 | 44,168 | 9,619,564 |
| Inferred HG005 hap1 | 2.76 G | 92.0% | 7,516,492 | 2,009,519 | 42,885 | 9,568,896 |
| Inferred HG005 hap2 | 2.76 G | 92.1% | 7,300,823 | 1,988,466 | 44,288 | 9,333,577 |

**Table S1. Benchmarking variant set statistics.** Benchmarking variant sets were generated based on 5 assemblies using a genome graph-based approach. Each set contains variants from 5 somatic samples in the HapMap mixture. SNV: single nucleotide variant; Indel: insertions and deletions < 50 base pairs; SV: insertions and deletions >= 50 base pairs. Covered Reliable Region: assembly reliable region defined by HPRC.

SNVs

| | All | Easy | Difficult | Extreme |
|---|---|---|---|---|
| GRCh38 | 0.9625; 0.7145 | 0.9920; 0.7548 | 0.9425; 0.7291 | 0.7908; 0.4802 |
| Pangenome (v3) | 0.9734; 0.7284 | 0.9927; 0.7673 | 0.9627; 0.7380 | 0.8506; 0.5069 |

Indels

| | All | Easy | Difficult | Extreme |
|---|---|---|---|---|
| GRCh38 | 0.7410; 0.1867 | 0.8914; 0.4489 | 0.7010; 0.1229 | 0.4573; 0.0863 |
| Pangenome (v3) | 0.7564; 0.1860 | 0.8963; 0.4470 | 0.7221; 0.1217 | 0.4781; 0.0864 |

SVs

| | All | Easy | Difficult | Extreme |
|---|---|---|---|---|
| GRCh38 | 0.6640; 0.2024 | 0.8881; 0.3519 | 0.8441; 0.2794 | 0.5309; 0.1579 |

| | | | | |
|---|---|---|---|---|
| Pangenome (v3) | 0.6598; 0.1939 | 0.8672; 0.3521 | 0.8452; 0.2805 | 0.5255; 0.1454 |

**Table S2. Performance (precision; recall) of the pangenome surjection approach using the HapMap mixture**. The performance was evaluated genome-wide and in easy, difficult, and extreme regions. The benchmarking was performed by comparing call sets with benchmarking variant sets based on GRCh38.

SNVs

| | All | Easy | Difficult | Extreme |
|---|---|---|---|---|
| GRCh38 | 0.9625; 0.7145 | 0.9920; 0.7548 | 0.9425; 0.7291 | 0.7908; 0.4802 |
| Inferred HG005 hap1 | 0.9764; 0.7819 | 0.9934; 0.8192 | 0.9649; 0.7857 | 0.8871; 0.5912 |
| Inferred HG005 hap2 | 0.9764; 0.7806 | 0.9934; 0.8183 | 0.9648; 0.7851 | 0.8865; 0.5861 |
| HG005 mat | 0.9691; 0.7801 | 0.9905; 0.8199 | 0.9612; 0.7860 | 0.8825; 0.5852 |
| HG005 pat | 0.9706; 0.7825 | 0.9906; 0.8223 | 0.9617; 0.7876 | 0.8853; 0.5895 |

Indels

| | All | Easy | Difficult | Extreme |
|---|---|---|---|---|
| GRCh38 | 0.7410; 0.1867 | 0.8914; 0.4489 | 0.7010; 0.1229 | 0.4573; 0.0863 |
| Inferred HG005 hap1 | 0.7650; 0.2585 | 0.9063; 0.4861 | 0.7384; 0.1810 | 0.5204; 0.1332 |
| Inferred HG005 hap2 | 0.7648; 0.2589 | 0.9068; 0.4854 | 0.7391; 0.1816 | 0.5194; 0.1339 |
| HG005 mat | 0.7548; 0.2563 | 0.8969; 0.4816 | 0.7343; 0.1833 | 0.5216; 0.1362 |
| HG005 pat | 0.7565; 0.2568 | 0.8969; 0.4823 | 0.7357; 0.1836 | 0.5227; 0.1366 |

SVs

| | All | Easy | Difficult | Extreme |
|---|---|---|---|---|
| GRCh38 | 0.6640; 0.2024 | 0.8881; 0.3519 | 0.8441; 0.2794 | 0.5309; 0.1579 |
| Inferred HG005 hap1 | 0.6642; 0.3202 | 0.8878; 0.4336 | 0.8499; 0.4152 | 0.5530; 0.2832 |
| Inferred HG005 hap2 | 0.6514; 0.3096 | 0.8711; 0.4075 | 0.8239; 0.4066 | 0.5454; 0.2767 |
| HG005 mat | 0.6950; 0.3108 | 0.9051; 0.4055 | 0.8865; 0.4037 | 0.6015; 0.2771 |
| HG005 pat | 0.7010; 0.3159 | 0.8959; 0.4253 | 0.8700; 0.4112 | 0.6043; 0.2818 |

**Table S3. Performance (precision; recall) of the pangenome-inferred DSA approach using the HapMap mixture**. The performance was evaluated genome-wide and in easy, difficult, and extreme regions. The benchmarking was performed by comparing each call set against its corresponding benchmarking variant set.

| Chrom | Location | Nucleotide change | Gene Name | Amino acid change | Type | AlphaMissense pathogenicity scores | Associated Cancer Types |
|---|---|---|---|---|---|---|---|
| Chr2 | 96031296 | G>A | GPAT2 | F72F | Synonymous | / | Breast cancer[3,4] |
| Chr2 | 110472682 | C>T | LIMS4 | E33K | Missense | 0.0835 | / |
| Chr2 | 112081091 | C>G | TMEM87B | I209M | Missense | 0.2065 | Melanoma[5] |
| Chr2 | 151589891 | C>T | NEB | D5082N | Missense | / | Renal cancer[6] |
| Chr6 | 28435526 | G>C | ZSCAN23 | Q164E | Missense | 0.1099 | Pancreatic ductal adenocarcinoma[7] |
| Chr7 | 144363604 | G>A | ARHGEF5 | R312K | Missense | 0.0889 | Colorectal cancer[8], Acute myeloid leukemia[9] |
| Chr8 | 7882517 | G>A | DEFB103A | Q51Q | Synonymous | / | Cervical cancer[10] |
| Chr9 | 35617156 | C>A | CD72 | G94G | Synonymous | / | Renal clear cell carcinoma[11], Acute leukemia[12] |
| Chr16 | 32253646 | G>A | TP53TG3D | R98Q | Missense | / | Breast cancer[13] |
| Chr18 | 13645428 | C>T | LDLRAD4 | P231L | Missense | 0.8379 | Colorectal cancer[14] |
| Chr19 | 30445460 | G>T | ZNF536 | C633F | Missense | 0.9983 | Neuroblastoma[15] |
| Chr19 | 39879573 | G>A | FCGBP | D2950D | Synonymous | / | Ovarian carcinoma, Lower-grade glioma etc. [16] |
| Chr19 | 40015580 | T>A | ZNF546 | V770V | Synonymous | / | Breast cancer[17] |
| hap#1#chr1#6 | 104980446 | G>A | NBPF1 | S7821F | Missense | / | Neuroblastoma[18], Adrenocortical carcinoma[19] |
| hap#1#chr22#5 | 18936680 | G>A | TRIOBP | R496Q | Missense | / | Gastric, rectal, pancreatic, and brain cancer[20] |
| hap#2#chr5#0 | 17607613 | C>T | TAF11L5 | K25K | Synonymous | / | / |
| hap#2#chr19#0 | 18879126 | C>T | FCGBP | P4615W | Missense | / | Ovarian carcinoma, Lower-grade glioma etc.[16] |

**Table S4. Summary of pangenome-inferred DSA unique SNVs located in exons.** hap#A#chrB#C indicates that the SNVs detected by pangenome-inferred DSA could not be lifted over to GRCh38. The amino acid change position for each SNV was determined based on the Ensembl canonical transcript. AlphaMissense[21] pathogenicity scores predict the likelihood of a missense genetic variant being pathogenic (0-0.34: likely benign; 0.34-0.564: uncertain; 0.564-1: likely pathogenic). hap#A#chrB#C: A: hap1/hap2; B: chromosome number; C: contig number.

variant effect prediction with AlphaMissense. Science *381*, eadg7492. https://doi.org/10.1126/science.adg7492.
